# Determining subpopulation methylation profiles from bisulfite sequencing data of heterogeneous samples using DXM

**DOI:** 10.1101/2020.10.12.333153

**Authors:** Jerry Fong, Jacob R. Gardner, Jared M. Andrews, Amanda F. Cashen, Jacqueline E. Payton, Kilian Q. Weinberger, John R. Edwards

**Affiliations:** Center for Pharmacogenomics, Department of Medicine, Washington University School of Medicine, St. Louis, MO, USA; Center for Data Science for Improved Decision Making, Department of Computer Science, Cornell University, Ithaca, NY, USA; Department of Pathology and Immunology, Washington University School of Medicine, St. Louis, MO, USA; Oncology Division, Department of Medicine, Washington University School of Medicine, St. Louis, MO, USA

## Abstract

Epigenetic changes, such as aberrant DNA methylation, contribute to cancer clonal expansion and disease progression. However, identifying subpopulation-level changes in a heterogeneous sample remains challenging. Thus, we have developed a computational approach, DXM, to deconvolve the methylation profiles of major allelic subpopulations from the bisulfite sequencing data of a heterogeneous sample. DXM does not require prior knowledge of the number of subpopulations or types of cells to expect. We benchmark DXM’s performance and demonstrate improvement over existing methods. We further experimentally validate DXM predicted allelic subpopulation-methylation profiles in four Diffuse Large B-Cell Lymphomas (DLBCLs). Lastly, as proof-of-concept, we apply DXM to a cohort of 31 DLBCLs and relate allelic subpopulation methylation profiles to relapse. We thus demonstrate that DXM can robustly find allelic subpopulation methylation profiles that may contribute to disease progression using bisulfite sequencing data of any heterogeneous sample.

## INTRODUCTION

DNA methylation changes have been implicated in a variety of human diseases, including cancer(1). Methylation measurements are typically made from heterogenous samples consisting of multiple cell-types, which each have unique methylation patterns. As such, it is frequently difficult to interpret whether an observed change in methylation is due to a shift in sample composition or due to a true change in the methylation state of an underlying cell-type. For example, tumors are comprised of both normal cell types and cancer subclones. These subclones can acquire changes that increase their fitness, leading to faster cancer progression, treatment resistance, and worse patient prognosis (2, 3). Though cancer subclones have generally been described with respect to genetic alterations, in principle epigenetic alterations such as DNA methylation could alter the expression of key genes in a subclone and impact its fitness. Moreover, in chronic lymphocytic leukemia (CLL), diffuse large B-cell lymphoma (DLBCL), and acute myeloid leukemia (AML), clonal heterogeneity in DNA methylation is associated with worse patient outcome(4) (5) (6). Unfortunately, even though subclonal methylation changes may be expected to underlie this observed heterogeneity, current methods do not effectively analyze subclonal methylation patterns from the bisulfite sequencing data of heterogeneous samples.

Ideally, we would like a computational approach to identify the number of underlying allelic subpopulations, their prevalence, and their respective methylation profiles from the methylation data of a heterogeneous input sample. Though extensive methods have been developed to describe subclonal architecture with respect to mutations or copy number variants (CNVs) (for a review see(7)), these methods cannot be easily adapted to methylation analysis since they typically assume subclonal events occur independently and are relatively rare. However, unlike mutations or CNVs, methylation changes often occur in blocks of multiple CpGs over a short range (differentially methylated regions, DMRs)(8, 9). Additionally, there are many more aberrant changes in methylation than mutations or CNVs. For example, frequently more than 100,000 DMRs are observed in solid tumors as compared to at most ~10,000 mutations or ~100 CNVs (10–12).

Most current analysis methods to determine the cellular composition and methylation profiles of subpopulation level events are developed for epigenome-wide association studies (EWAS) (13–17). However, EWAS uses array-based technologies that probe the methylation state of 3% of the CpGs in the human genome, and as such, these approaches use assumptions and error-models that are not appropriate for sequence data. For example, they do not consider the strong local correlations of methylation changes, since adjacent probes are frequently greater than 1 kb apart(18).

Thus, we have developed DXM (<u>**D**</u>econvolution of Subpopulations E<u>**x**</u>isting in <u>**M**</u>ethylation Data), a novel deconvolution strategy to identify the major allelic subpopulations, their relative prevalence, and their respective methylation profiles from a heterogeneous sample (Fig. 1a,b). DXM does not require explicit prior knowledge of the number of subpopulations or what types of cells to expect, and it provides a framework for considering methylation differences across multiple CpGs at the single-CpG resolution offered by bisulfite sequencing data. We benchmarked DXM on a wide set of simulated mixtures using bisulfite sequencing reads from sorted hematopoietic cell types and found that DXM outperformed methylPurify(19), another method developed for subclonal analysis of bisulfite sequencing data. We further conducted Agilent Methyl-Seq bisulfite sequencing analysis in four DLBCLs and validated that DXM predictions for subpopulation methylation profiles were recapitulated in relevant sorted CD4^+^ T and CD19^+^ B cells. As proof-of-concept, we applied DXM to a cohort of 31 DLBCLs(5) to highlight how DXM can be used to analyze subpopulation methylation in heterogeneous cancer samples and relate them to relapse.

**Figure 1.**
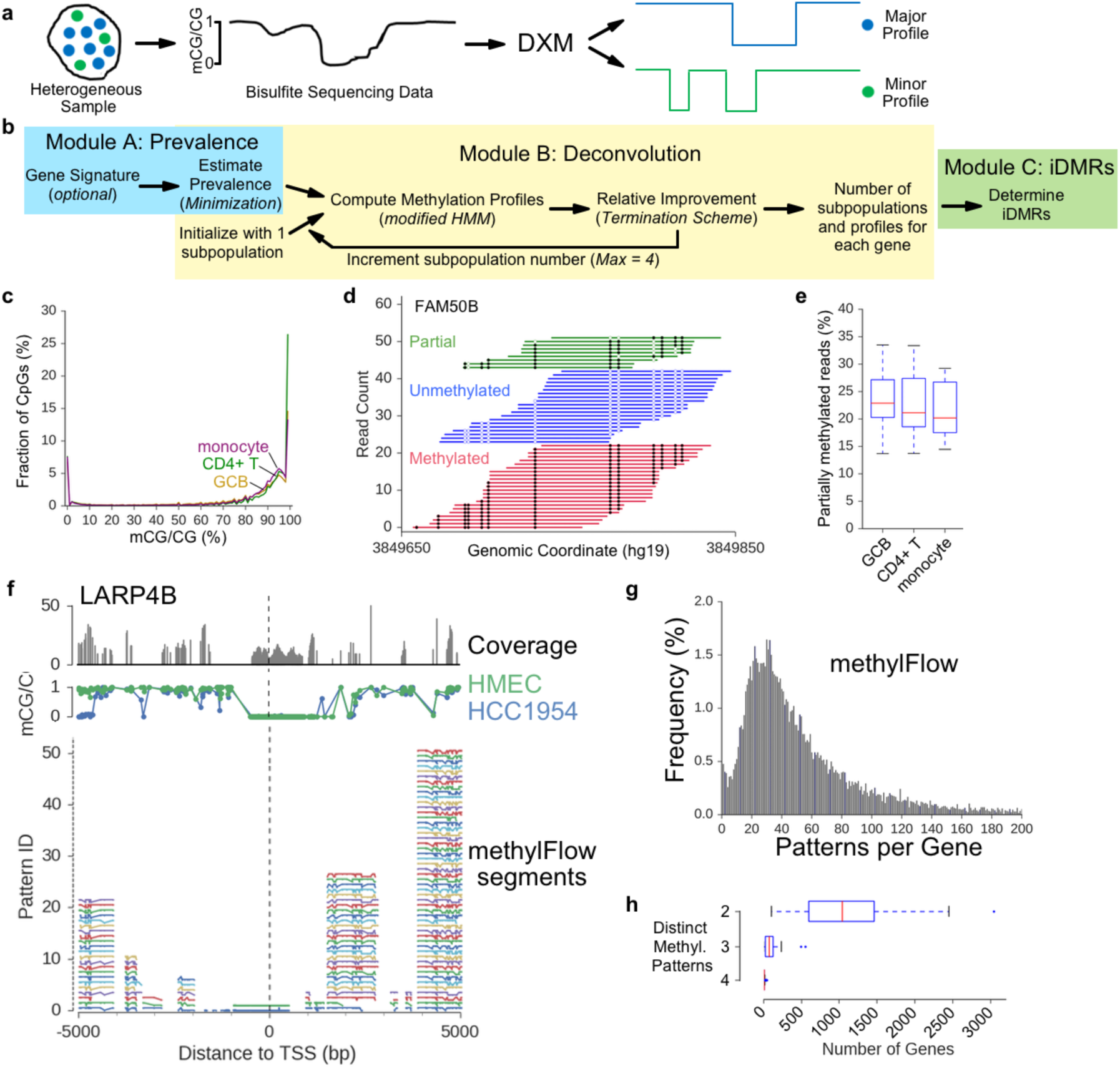
DXM scheme and design considerations. **a)** DXM takes bisulfite sequencing data from a heterogeneous sample and identifies its underlying allelic subpopulation methylation profiles, the number of subpopulations, and their relative prevalence. **b)** DXM consists of three modules: Module A estimates the prevalence, Module B solves subpopulation methylation profiles, and Module C calls intrasample DMRs (i-DMRs). **c)** Fractional methylation distribution for sorted cell types. 12.4%, 15.5%, and 22.7% of CpGs are partially methylated (between 20-80% methylation) in CD4^+^ T-cells (green), GCB cells (gold), and monocytes (purple), respectively. **d)** Bisulfite sequencing reads across part of the imprinting control region for FAM50B in germinal center B-cells (GCBs). Open circles are unmethylated CpGs and closed circles are methylated. **e)** Number of partially methylated reads at 19 well-characterized imprinted loci in different sorted cell types. **f**) methylFlow (MF) outputs for LARP4B from analysis of a 22x coverage HMEC-HCC1954 mixture (35:65). For individual segments, MF outputs between 1 and 52 potential profiles. **g)** The most profiles in any individual segment by MF for all promoters in a 22x coverage HMEC-HCC1954 mixture (35:65). **h)** Number of gene promoters with DMRs identified between or among different sorted cell types (CD4T, CD8T, erythroblast, eosinophil, hematopoietic multiprogenitor, GCB, megakaryocyte, monocyte, osteoclast). Promoters were considered to have distinct methylation patterns if there was a DMR identified between each pair of cells considered (e.g. for cell types A, B, and C, there are three distinct patterns if there is a DMR between A-B, B-C, and A-C). 17,450 total genes were considered.

## MATERIALS AND METHODS

### Public Datasets

Bisulfite sequencing data was downloaded from the Roadmap Epigenomics project(20) (REP), the Blueprint Epigenome Project(21) (BEP), ENCODE(22), and GEO (Accession: GSE29069(23), GSE66329(24)). A full list is provided (Supplementary_File_1.xlsx).

### DXM

A scheme for DXM is provided in Figure 1a-b. DXM takes methylation data from heterogeneous samples as input and deconvolves it into the number of major underlying allelic subpopulations, their prevalence, and their respective methylation profiles. DXM uses processed data as input (BED-like format: tab-delimited columns of chromosomes, start_position, end_position, methylation_level, sequencing_coverage, region_id), consisting of the average methylation mCG/CG and sequencing coverage at each CpG site and performs deconvolution over user-specified intervals (e.g. promoter regions or enhancers) without the need for a specific coverage cutoff. Typically, we recommend collapsing CpG data across strands to increase reliability of the methylation estimates, but this is not required (pre-processing details for specific experiments are detailed below). We adopt an iterative scheme to solve for the number of subpopulations, beginning with only 1 major subpopulation, and adding additional subpopulations provided that they do not cause an overfit. This approach does not assume how many underlying subpopulations are present *a priori*, but it takes an Occam’s Razor approach in determining the minimal number of subpopulations that can reasonably explain the observed data.

#### Minimization

We next estimate the best possible prevalence of subpopulations from the distribution of all fractional methylation values detected. For example, if there are two subpopulations with prevalence of 0.3 or 0.7 in the sample, then, given that each subpopulation will be methylated or not, the expected fractional methylation values detected are {0, 0.3, 0.7, 1}. Thus, we minimize the L1-difference between the set of all combinations of underlying fractional methylation values with the original methylation distribution. In the (unlikely) event of a tie, the solution with the smallest possible subpopulation is selected. Each gene’s methylation profile is then solved independently as follows.

#### Modified Hidden Markov Model

Given the number of underlying subpopulations and their expected prevalence, we then solve for the most likely methylation profiles by applying the Viterbi algorithm to a modified Hidden Markov Model (HMM). An HMM is well-suited to model a sequence of data with local correlations, as exhibited by DNA methylation, and is characterized by its states, transitions, and emissions(25). For our modified HMM, if there are *d* subpopulations to consider, a state is a length-*d* vector of the binary methylation values of each underlying subpopulation at a given CpG. The transition probability represents the change in methylation of each underlying subpopulation from one CpG to an adjacent CpG. To calculate the transition probability, we have defined a set of 2×2 transition matrices that capture transitions of one subpopulation’s methylation state from one CpG to the next. From this set, the transition matrix is selected based on the distance between CpGs, which captures correlations in methylation and distance seen in Supplemental Figure 2. We consider each subpopulation’s transition to be independent and identically distributed, so the transition probability is ultimately calculated from *d* identical transition matrices.

**Figure 2.**
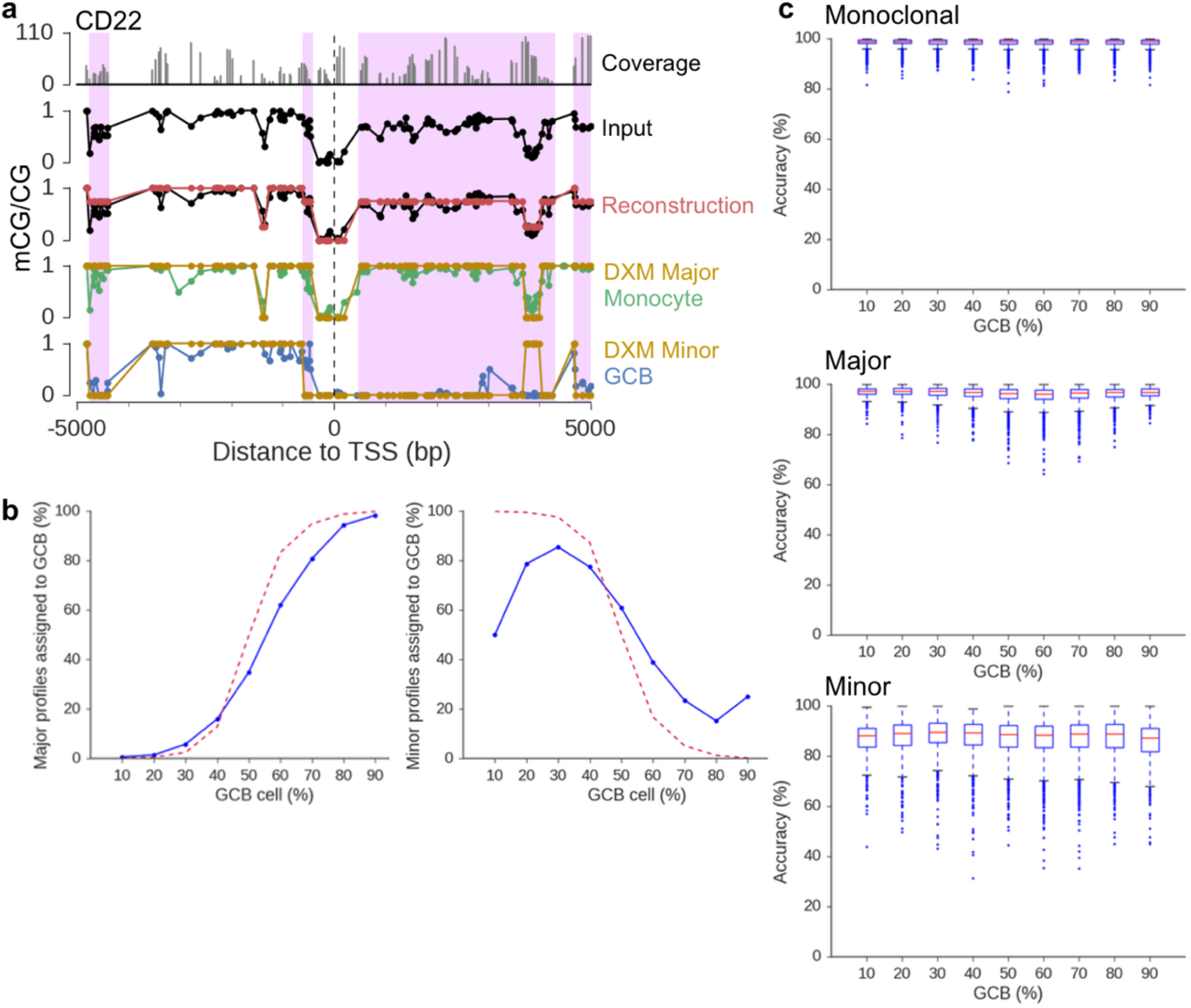
DXM solves for subpopulation methylation profiles and their prevalence in heterogeneous mixtures. **a)** DXM solution for CD22 for a 55x coverage simulated mixture of monocytes and GCB (35:65). **b)** Number of gene promoter profiles assigned to GCB cells (blue) across 55x coverage mixture simulations with different GCB prevalences. The dotted red line indicates the number of CpGs with more subsampled reads from GCB cells than monocytes. **c)** Accuracy of DXM methylation profiles with respect to reference GCB or monocyte profiles for promoters where DXM identified the same number of methylation profiles as expected by DSS.

To ensure rigor, robustness, and reproducibility, transition probabilities were trained on somatic tissue and cell-line datasets available via REP, and all testing was conducted on datasets of different cell types available for BEP or GSE29069. Briefly, for all CpGs in a dataset, the detected fractional methylation was rounded to the closest binary methylation state (fully unmethylated or fully methylated). Transitions between binary methylation states of adjacent CpGs were then counted (empirical sampling). We elected to group all transitions greater than 1000bp in the same category when selecting transition probabilities to use; this case is infrequent in the TSS±5kb region. We do not utilize cell type-specific transition probabilities for underlying subpopulations.

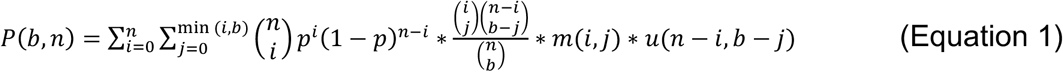

The emission probabilities, *P(b,n)*, were calculated as follows (Equation 1). There are *n* reads, *b* of which are methylated at a given CpG. Assuming reads are independently and identically distributed among allelic subpopulations as a function of prevalence, reads can be grouped by whether they came from an allelic subpopulation whose underlying state was methylated or not. Thus, if *i* reads came from subpopulations with an underlying state of unmethylated, *n-i* come from those with an underlying state as methylated, and this case follows a binomial distribution, where *p* is the prevalence of all subpopulations with an underlying methylated state. Of these *i* reads, *j* may be observed as methylated, and assuming all reads are independent, the probability of this event follows a hypergeometric distribution. Lastly, the probability that *j* of *i* reads are observed as methylated, given that the expected underlying state is fully methylated, is modeled with a beta-binomial distribution *m*. A separate beta-binomial distribution, *u*, models a similar probability that *b-j* of *n-i* reads are observed as methylated given an expected underlying state of fully unmethylated. Determination of the beta-priors for these beta-binomial distributions is provided below (see below in *Beta-binomial fit*). Summing across all valid combinations of *i,j* gives the final emission probabilities. We use log-probabilities in all calculations to ensure precision.

#### Termination Scheme

DXM utilizes a termination scheme to determine if an additional allelic subpopulation better explains the observed data. Let *p*(*x*) represent the probability that the state-sequence *x* correctly explains the data, and let *x*_*r*_ represent the most likely state-sequence for *r* subpopulations. DXM will consider an additional subpopulation to be valid if there is an increase in likelihood, that is if *p*(*x* = *x*_*r*+1_) > *p*(*x* = *x*_*r*_). By Bayes’ rule, this can be written as *p*(*x* = *x*_*r*+1_|*r* + 1) ∗ *p*(*r* + 1) > *p*(*x* = *x*_*r*_|*r*) ∗ *p*(*r*), where *p*(*r*) is the probability that *r* subpopulations best explain the underlying data. In the uninformed case, we would have a uniform prior on the number of subpopulations to expect, e.g. *p*(*r*) = *p*(*r* + 1), and thus, the comparison is *p*(*x* = *x*_*r*+1_|*r* + 1) > *p*(*x* = *x*_*r*_|*r*). The conditional probability of a state-sequence *xr* explaining the data given *r* subpopulations, *p*(*x* = *x*_*r*_|*r*), is solved by the Viterbi algorithm, so termination is achieved when the relative probability of a solution is worse. Though it is possible that our scheme does not yield the global solution {e.g. *p*(*x* = *x*_*r*+1_|*r* + 1) ≤ *p*(*x* = *x*_*r*_|*r*) but *p*(*x* = *x*_*r*+2_|*r* + 2) > *p*(*x* = *x*_*r*_|*r*)}, we have not observed this in practice and believe the case to be highly unlikely for reasonably well-behaved beta-binomial distributions. Additionally, this scheme is guaranteed to converge. Consider the case where an additional subpopulation has the same methylation profile as an existing one. In this case, the emission probability would remain unchanged, but the transition probabilities would be less likely since there is an additional subpopulation to consider. Since there are a finite number of possible methylation profiles for *g* CpGs (2^g^), this case is guaranteed, though in practice we need to consider far fewer than 2^g^ methylation profiles (e.g. 4 total profiles).

#### DXM output

The output of DXM includes an estimate to the number of underlying subpopulations, their respective methylation profiles, and an estimate of their relative prevalence in the sample. Given the utility of identifying differentially methylated regions (DMRs), we have also provided a post-hoc utility to identify intrasample DMRs between or among identified subpopulation methylation profiles (i-DMRs). The DMR definition was adapted from DSS (>50bp, >3CpGs, >90% CpGs are differentially methylated). DXM i-DMRs can be ranked based on their improvement in relative probability when modeled with 2 subpopulations as compared to 1, given as *p*(*x* = *x*_2_|2) − *p*(*x* = *x*_1_|1), which provides compatibility with DXM results for analyses such as gene set enrichment.

### Beta-Binomial Fit

An empirical beta-prior for use in DXM was drawn from training data from REP. For an expected unmethylated state, every CpG with percent methylation between 0% and 40% was considered; similarly, 60%-100% were the cutoffs for an expected methylated state. The methylation of each CpG was rounded to the nearest 0.5%, and these discrete empirical distributions were then normalized. To find a suitable beta prior for this distribution, the probability density function of a given beta distribution was evaluated at each domain value in the empirical distribution (e.g. every 0.5% methylation). This new distribution was then normalized, and the L1-difference between it and the empirical distribution was minimized. All alpha/beta parameters for underlying beta distributions were fixed as integers, and a linear combination of beta distributions was employed to better capture spiking behavior near the edges of the distribution (e.g. 0% or 100% methylation) and a small signal around 50% methylation.

### Promoter region definition

For analysis, genes were defined with respect to RefSeq(26) annotations, where only genes (transcripts) with the ‘cmpl’ annotations for both the coding start and end sites were used. For genes with multiple transcription start sites (TSS), only the first was considered. Only genes on autosomal chromosomes were considered for analysis. Gene promoter regions were defined as the region +/− 5kb from the annotated TSS. While there is no agreed upon definition of a gene promoter, this region was chosen because it is larger than available definitions, and this window has previously been shown to be useful for predicting expression changes (27).

### Methylation-level Mixture Simulations

Average methylation levels (mCG/CG) within 5kb of the TSS for each cell-type were rounded to 0 or 1 to binarize profiles for individual samples. For each simulation at a fixed prevalence and a fixed coverage, 1000 genes were randomly selected. The mixed methylation level (mCG/CG) of each CpG was then computed as the dot product of the prevalences and methylation values. Binomial sampling was then used to simulate the measured methylation value of the mixed sample based on the chosen coverage. DXM was then used to deconvolve the mixture.

### Read-level Mixture Simulations

For simulated mixture generation, bisulfite sequencing reads (from BEP or GSE29069 datasets) were mapped with bsmap2.90(28) against the hg19 genome using the -R flag. Mapped bisulfite sequencing reads were coordinate-sorted with tabix(29), after which all reads overlapping the TSS±10kB window of genes were taken for input. Next, reads were subsampled with two parameters: the expected average coverage of the total mixture, and the expected prevalence of each underlying cell type. Given an expected average coverage, the expected total number of reads was calculated as (coverage * *domain size* / read length). Average read length was 100bp, and domain size was estimated as the number of genes * 10kb (the size of the TSS±5kb window). Next, the expected number of reads from each cell type was calculated as prevalence * total reads. Given the number of reads from each cell type, we calculated a sampling rate for each file, assuming single-end reads are present. This setup allows us to sample differences in coverage across each region and does not impose that more reads must come from the more prevalent cell type, which may not be true upon sequencing. Once a simulated mixture was generated, its processed methylation was obtained using the “methRatio.py” script provided with bsmap, and methylation data from both strands of DNA were consolidated for analysis.

### DMR Calls

We utilized DSS(30) as an independent DMR-caller to help evaluate DXM results. DSS was run on each sample using the following parameter set: equal dispersion = True (for comparing single samples against each other), deltaVal = 0.3 (“30% methylation difference”), and a pval threshold of 0.05 (a CpG is considered differentially methylated if this threshold is met).

### Gene Signatures for Prevalence Calling

Gene signatures for prevalence calling were defined by running DSS to identify DMRs between GCB (Blueprint: T14_11) and monocytes (Blueprint: S000RD).

### MethylPurify and MethylFlow

MethylPurify(19) and methylFlow(31) were run with default settings. For methylPurify, the most informative bins were identified from the “*Informative_bins.bed.OneForCGI” file, and subpopulation profiles were found in the “MethylProfile.bed” file. For methylFlow, methylation of segments were obtained from the “patterns.tsv” file, where pattern id (pid) refers to the number of profiles solved for a segment.

### Evaluation Metrics

To evaluate whether DXM-solved methylation profiles resembled those of reference cell-types, we first computed the “closest possible” reference methylation profile by rounding all fractional methylation values of the reference cell types to either 0 (fully unmethylated) or 1 (fully methylated). DXM-solved methylation profiles were then compared against each closest possible reference methylation profile and assigned to the reference that differed at the fewest number of CpGs (e.g. L1-norm difference). During assignment, DXM-solved subpopulations were not forced to be associated to different reference cell types, and assignment was allowed even in cases where there was significant discrepancy (e.g. >50% of CpGs).

Accuracy was reported as the percent of CpGs that were incorrect out of all CpGs for that gene promoter. After each profile was assigned to a reference cell type, to evaluate how accurate DXM was in identifying cell types at the correct relative prevalence, the major allelic subpopulation was defined from the expected prevalence of the underlying subpopulations in the mixture. The percent of major subpopulation profiles DXM identified that corresponded to the correct expected major cell type was then computed.

### Ontology Analysis

Ontology analysis was conducted using the DAVID functional annotation tool with default parameters and background as *H. sapiens*.

### Primary samples

Primary DLBCL samples were obtained from the WUSM Lymphoma Banking Program. Written informed consent was obtained from all patients as part of the WUSM Lymphoma Banking Program. This study was approved by the Washington University in St. Louis Institutional Review Board (#201710120). A full table of sample characteristics is provided in (Supplemental Table 2) All samples were cryopreserved at >10 million cells/ml. Samples were flash-thawed at 37°C and pelleted at 200 rpm, 5 min, 4°C. Samples were then resuspended in wash buffer (4% Fetal Calf Serum in dPBS). Half of each sample was isolated for Agilent-Methyl Seq, and the remaining half of each sample was sent for cell sorting.

### Cell Sorting (FACS)

Each sample was first blocked with 100ul of 5% Human TruStain FcX (Biolegend) in wash buffer (4% FCS in dPBS) for 7min at RT. Samples were pelleted (200g, 5 min, 4°C) and resuspended in 100ul of the following binding buffer: 20ul (1 test volume) mouse anti-human CD19-PE (BD Biosciences), 20ul (1 test volume) mouse anti-human CD4-FITC (BD Biosciences), 5ul (1 test volume) 7-AAD (BD Biosciences), and 45ul wash buffer. Samples were incubated in the dark at 4°C for 20 min. Following incubation, samples were washed twice with wash buffer before sorting with BD FACSAriaII at a standard flow rate for PE, FITC, and PerCP-Cy5.5-A channels. CD4+/CD19-/7-AAD-cells and CD19^+^/CD4-/7-AAD-cells were collected for subsequent analysis.

### Agilent Methyl-Seq Analysis

Genomic DNA was isolated from each sample using the Zymo Quick-DNA Miniprep kit. Agilent Methyl-Seq was then conducted for each sample following manufacturer’s recommendations for 1ug of input gDNA. Bisulfite conversion was conducted with Zymo Methylation-Gold kit, and samples were sequenced with an Illumina Hi-Seq3000 (2×150 paired-end reads). Sequencing specifications are found in a table. On average, we obtained >50x coverage across ~4.9 million CpGs for each patient, and we observed a minimum bisulfite conversion efficiency of >98.9% for all samples, as estimated by nonCpG conversion rate. All data were then aligned with biscuit v0.3.8 (https://github.com/zwdzwd/biscuit) and processed methylation was obtained with the BISCUIT pipeline.

### Targeted Bisulfite Sequencing

A table of all bisulfite sequencing primers and locations is provided in Supplemental Table 5. Bisulfite primers for loci of interest were designed using methPrimer2.0(32) with default settings and BiSearch(33), requiring only one major genomic location for expected amplification. Primers were temperature-optimized for PCR amplification (QIAGEN Pyromark PCR kit) on bisulfite converted gDNA from GM12878 cells. Each sample had gDNA isolated and was bisulfite converted as above. PCR-amplification was conducted for each region in each sample using 100pg of bisulfite-converted DNA as input. PCR products were barcoded and made into Illumina sequencing libraries as in (34). Libraries were sequenced on an Illumina Hiseq 4000. Sequences were trimmed for adaptors and poor-quality sequence using TrimGalore with default parameters. Sequences were then mapped to their target regions using Bismark (35) and default parameters and methylation levels (mCG/CG) extracted using bismark_methylation_extractor.

### Statistics

Distributions were compared using Student’s t-test for two groups or ANOVA with Tukey’s posthoc t-test for multiple groups. Cohen’s d-test was used to estimate effect size. Enrichment of CpGs in genomic elements (CpG island, shore, other elements) was compared with a Chi-square test for proportions relative to either all CpGs in the +/− 5kb region around the TSS or to CpGs covered by Agilent Methyl-Seq.

## RESULTS

### Features of Methylation Distributions in Sorted Cells

Before developing a deconvolution strategy, we sought to understand what constitutes distinct DNA methylation profiles in bisulfite-sequencing data. We first examined the distribution of methylation levels (mCG/CG) within individual sorted cell types. One would expect that within a sorted cell type there would be peaks at 0%, 50%, and 100% corresponding to unmethylated, imprinted, and methylated CpGs, with some noise. However, from inspecting the distribution of methylation values (Figure 1e), there are a variety of intermediate methylation values (i.e. not fully methylated or unmethylated) as well as substantial shoulders to the expected peaks around 0% and 100% methylation.

We next examined methylation in three hematological cell types at imprinted loci. Imprinted loci are characterized by a DMR that is fully methylated in one inherited allele and fully unmethylated in the other(36, 37). Despite this, we find that 20% of bisulfite sequencing reads across DMRs in three hematological cells show partial methylation, or evidence for methylated and unmethylated CpGs on the same allele (Figure 1c-d). This results in a fractional methylation distribution at imprinted DMRs that is wider than what might be expected for a profile with 50% methylation (one methylated and one unmethylated allele, Supplemental Figure 1).

These deviations from idealized distributions at imprinted domains, and fully methylated and unmethylated CpG sites likely do not arise from technical issues since both sequencing read quality (Supplemental Figure 1) and bisulfite conversion efficiency are high (>99.7%) (38). As such, these likely represent noise due to further biological heterogeneity or biological noise such as age-associated drift (39) that is frequently smoothed to facilitate biological interpretation. Taken together, a binomial distribution, defined by number of reads and expected average methylation, is insufficient to model these data. We will thus incorporate a beta-binomial distribution, whose parameters are learned from real-sequencing data to incorporate these phenomena.

### Considering every methylation change at each CpG in every sequencing read leads to an intractable solution

We next applied methylFlow (MF)(31) to simulated mixtures of reads from whole-genome bisulfite sequencing of HMEC (Human Mammary Epithelial Cells) and HCC1954 (breast tumor) cells to understand whether interpreting every methylation change at each CpG as an individual pattern could lead to a useful result. methylFlow uses a network flow analysis to stitch together segments of methylation patterns and enumerate all possible methylation profiles supported by bisulfite sequencing reads from an experiment. From a 22x coverage simulated mixture of HMEC:HCC1954 (35:65) methylFlow identified 17,367 of 17,450 (99.5%) gene promoters (TSS +/− 5kb) as having multiple methylation profiles (Figure 1f). For comparison, the DMR-caller DSS identifies 1,203 of 17,450 promoters with differential methylation. Within each promoter window, in the segment with the most potential profiles, methylFlow found on average ~55.7 profiles (Figure 1g). However, the methylFlow segments are relatively small with a median length of 200-750 bp(31), thus enumerating the potential combinations of these profiles yields many more potential profiles at each promoter. As an example, there are upwards of 1.7*109 possible promoter profiles from combining predicted segments for LARP4B across the 10 kb region centered at the TSS (Figure 1f). While we can’t rule out that each of these possibilities is real and not due to biological and technical noise, in practice this result is difficult to interpret and leads to an intractable solution. Similar results were obtained for simulated mixtures of GCBs and monocytes (data not shown).

This result suggests it would be better to consider regions of changes in methylation rather than every individual change at a CpG site. In fact, the functional significance of changing DNA methylation at a given CpG is unclear. DNA methylation levels are highly correlated over 100s to 1000s of bases (Supplemental Figure 2)(8, 9), and methylation changes frequently co-occur as cell type-, tumor-, allele-specific (imprinted) DMRs. Further, the best associations between expression and methylation are found considering the entirety of methylation changes around the promoter(25).

### Number of cell type-specific promoter methylation patterns

An important consideration in any deconvolution strategy is how many different methylation patterns are expected in a complex mixture. To understand how many different methylation patterns between cell-types would be expected in mixtures of hematologic cell types we called DMRs between different cell types and examine how many unique patterns existed across the different cell types. We find that for a given promoter, there are rarely, if ever, four or more distinct DMRs between cell types in a given mixture (Figure 1h).

Based on these observations, we developed DXM to determine subpopulation methylation profiles spanning a pre-defined set of genomic regions (e.g. a +/− 5kb promoter window for expressed genes), as well as a prevalence estimate for each allele, and a list of intrasample differentially methylated regions (i-DMRs), or regions where allelic subpopulations have differential methylation. DXM considers regions of change rather than individual changes, effectively smoothing over individual changes.

### DXM

DXM considers each region (e.g. promoter window) separately, reflecting how two distinct allelic subpopulations do not need to have distinct methylation patterns at every region. For each region, DXM uses a modified heterogeneous factorial Hidden Markov Model (HMM, Fig. 1b). The transition probabilities of the HMM are used to capture the local correlations between methylation states at nearby CpG sites. The emission probabilities of the HMM incorporate a beta-binomial distribution to empirically model and smooth over noise in the bisulfite sequencing data. Lastly, DXM determines the number of allelic subpopulations iteratively, beginning with one subpopulation and progressively considering additional subpopulations to determine an optimal solution. The resulting methylation profiles solved by DXM are binarized, reflecting how for a given strand of DNA, at a CpG, a methyl group is either present or absent at the 5 position of cytosine.

### DXM performance on simulated idealized mixtures

We first demonstrated DXM performance in simulated idealized cases. To remove potential noise, methylation levels (mCG/CG) were binarized to 0 and 1 for each cell type. For a fixed coverage, unmethylated and methylated counts for each CpG site were modeled using a binomial distribution. This effectively assumes that errors from inadequate bisulfite conversion and sequencing are negligible relative to the binomial sampling error (see methods for details). As seen in these initial simulations, DXM performs very well. The accuracy of each profile is very high across most coverages (see methods for details), while the number of genes with multiple profiles increases with coverage until ~30-90x depending on the prevalences of the underlying cell-types (Supplemental Figure S3, S4). This demonstrates that DXM is generally optimized to only call profiles when it can do so accurately, otherwise is refrains.

### DXM accurately solves subpopulation methylation profiles in heterogeneous mixtures

To determine the effectiveness of DXM, we generated a series of simulated mixtures by subsampling bisulfite sequencing reads from sorted germinal center B-cells (GCB, from tonsil(21)) and monocytes (from cord blood(21)) at a fixed expected average coverage of 55x. Our simulations included cases where there were more GCB as well as more monocytes. For our studies, we have initially considered methylation profiles in a ±5kb window around the transcription start site (TSS). This is a conservative estimate of the promoter region, which includes the regions where differential methylation most likely impacts gene expression(40). We then applied DXM to deconvolve these mixtures (example output in Figure 2a, Supplemental Figure 5). To quantify DXM performance, we first identified gene promoter windows expected to have differentially methylated regions (DMR) between cell types using DSS(30), a DMR-caller that uses a Bayesian beta-binomial hierarchical model.

We examined the degree of concordance between DXM i-DMRs predicted from simulated cell mixtures and DMRs identified from the underlying cell types. DXM on average recapitulated about 64% of all DMRs between the underlying subpopulations (Supplemental Figure 6a). Across all simulated mixture ratios, we found that DXM predicted a much larger number of i-DMRs for GCB-monocyte mixtures relative to the number of DMRs DSS identified between each underlying cell type (e.g. for a 30:70 mixture, 19,755 i-DMRs and 3,012 DMRs, or 5,722 vs 2,187 gene promoters). The average coverage, length, number of CpGs, and methylation levels of partially methylated CpGs for i-DMRs and DMRs are similar (Supplemental Figure 7).

We next evaluated the accuracy of deconvolved promoter profiles output by DXM for genes where DMR-analysis agreed with the number of DXM predicted allelic subpopulation profiles. For each promoter, DXM solved for subpopulation methylation profiles, which were then assigned to the closest reference cell type. As expected, the percent of profiles assigned to the GCB cell type in the major subpopulation increases as the number of CpGs for which more reads are subsampled from the GCB cells is increased (Figure 2b). The inverse relationship can be seen for the minor subpopulation. DXM performs best when the minor subpopulation comprises 20% or more of the mixture. This might be expected for our simulations, since in a 55x coverage mixture, a 10% subpopulation corresponds on average to only 5.5 reads. For some promoters, especially when the prevalence of the minor subpopulation is less than 20%, the minor methylation profiles output by DXM are closer to the true major subpopulation reference profile, rather than the minor population reference (Supplementary Figure 8). This observation likely reflects how intrinsic variation in typical methylation data from single sorted cell-types at fully methylated and unmethylated CpG sites masks the ability of deconvolution approaches to detect low prevalence subpopulations (Fig. 1c-d).

DXM achieved high accuracy, correctly identifying methylation across 98.5% of CpGs for promoters with one methylation pattern, 96.2% for major methylation profiles, and 87.1% for minor methylation profiles (average performance across all prevalence mixtures for GCB-monocytes, Figure 2c). Similar results were obtained for mixtures generated from sorted CD4^+^T-CD8+T cells (Supplemental Figure 9) as well as from a mixture of non-tumorigenic human mammary epithelial cells (HMEC) and a breast cancer cell line (HCC1954, Supplemental Figure 10). Increasing sequencing coverage up to 88x did not impact reconstruction accuracy (Supplemental Figure 11), consistent with earlier simulations (Supplemental Figures 3 and 4).

The DMRs that were not recapitulated by DXM tended to be shorter and have fewer CpGs (Supplemental Figure 7). Additionally, we found that increasing sequencing coverage did not improve this overlap, though it increased the total number of i-DMRs and DMRs found (Supplemental Figure 6b). Thus, we suspected that this difference was due to the domain of input data we considered. We next isolated each “missed” DMR and reran DXM for just that DMR instead of across the full +/− 5kb promoter window, and we found that on average, DXM recapitulated ~86% of these “missed DMRs” (Supplemental Figure 6c). Taken together, this highlights how DXM and DSS use different approaches to smoothing the data and locating DMRs, but can recapitulate similar results if forced to.

### DXM identifies substantial subpopulation methylation profiles in sorted cell types

We next sought to understand why DXM predicted many more iDMRs than expected based on DMR calling between the cell types. Plotting the distribution of methylation levels (mCG/CG) for the reference GCB and monocytes shows that these levels are often similar between cell-types but in an intermediate methylation state (e.g. 30% methylation in GCB and 30% in monocytes) (Figure 3a). One interpretation is that for these loci, there are distinct DNA methylation patterns in subpopulations, but these subpopulations differ from the expected reference cell types as might be defined from sorting. Supporting this is the observation that when we applied DXM to only GCB cells or only monocytes alone, we found that 70% of the DXM-specific i-DMRs predicted in the GCB:monocyte mixture were also detected as i-DMRs within only GCB cells or only monocytes. However, it is unlikely that the multiple patterns found within these cell types represent what is traditionally referred to as biological heterogeneity (e.g. multiple cell subtypes within a sorted cell type), since the sub-profiles are mostly shared by two distinct sorted cell-types: GCBs and monocytes. For instance, when we look at i-DMRs that lie within nine total sorted cell types (CD4+ T, CD8+ T, eosinophil, erythroblast, GCB, mematopoietic multipotent multiprogenitor, megakaryocyte, monocyte, osteoclasts), we detected 840 promoters with an i-DMR within all nine cell-types and ~500 genes where five cell-types exhibited the i-DMR (Figure 3b). The distribution of methylation levels of CpGs in these common i-DMRs is not centered around 50%, suggesting that they are likely not imprinted (Figure 3c).

**Figure 3.**
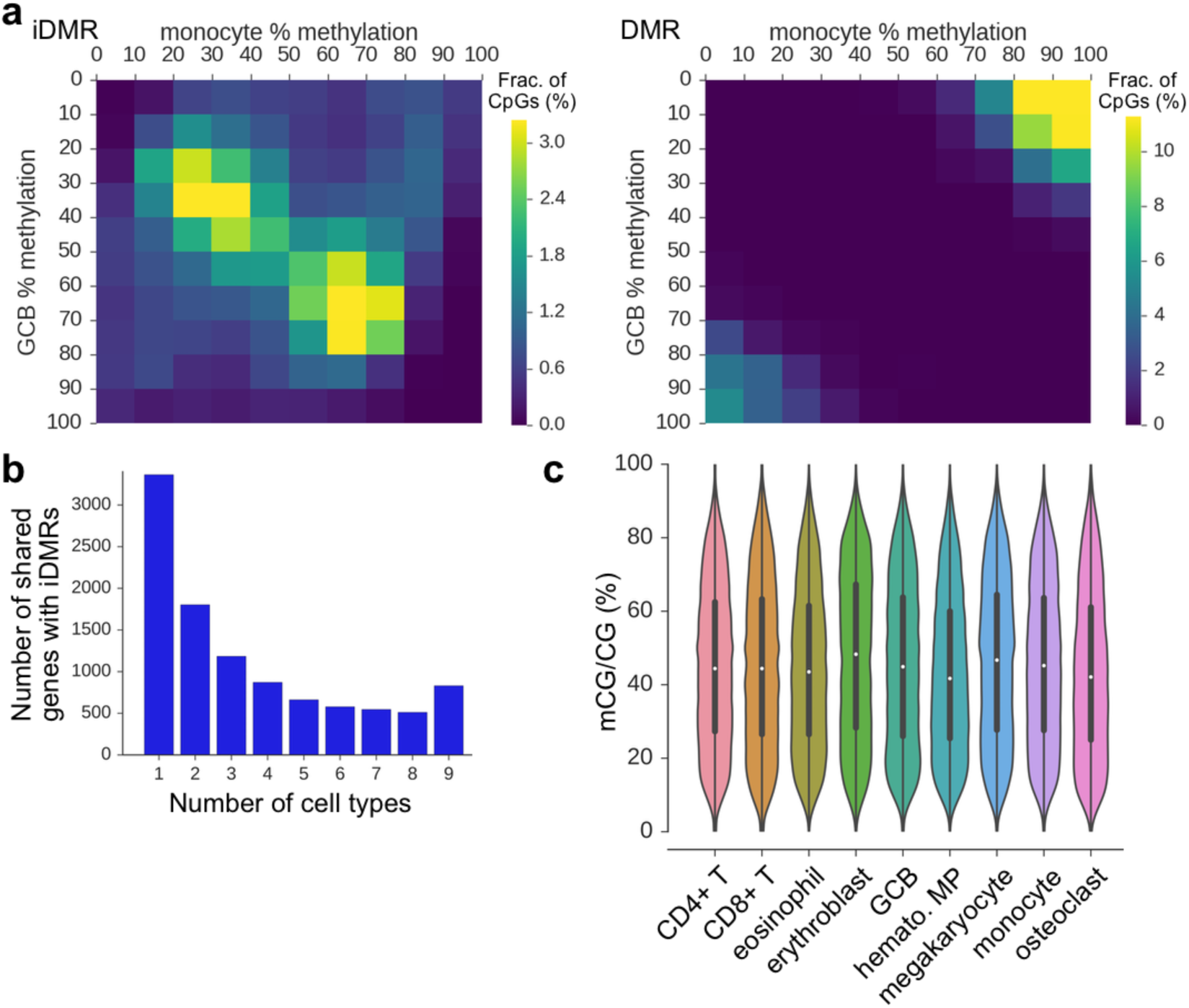
DXM identifies many i-DMRs within sorted cell types. **a)** i-DMRs and DMRs identified in a 55x mixture of GCB:monocytes (30:70) have different methylation profiles of the underlying reference GCB cells and monocytes. For instance, many i-DMRs are found with 30-40% methylation in both GCB and monocytes. **b)** The number of gene promoters with i-DMRs that are shared between different numbers of sorted cell types is shown. There are substantial numbers of i-DMRs that are both unique to individual cell-types, as well as shared across many cell types. The cell types used are listed in (c). **c)** Fractional methylation distribution of CpGs in i-DMRs for sorted cell types.

Ontology analysis indicates that i-DMRs that are in shared across all 9 hematologic cell-types are enriched for genes associated with cadherin-domain proteins (Supplemental Table 1). Cadherins are critical to cell-adhesion(41), which regulates many aspects of leukocyte function, including extravasation and vascular permeability for circulating leukocytes(42). This suggests there is a methylation signature across these genes associated with reduced cadherin expression and thus decreased tendency of a particular cell to adhere (e.g. make it less sticky). Further, because cell-adhesion is a pathway common to all cells, this is consistent with the finding that there are multiple methylation profiles within sorted cell types that may not correspond to traditional cell subtypes but certainly could represent distinct biological states. Thus, DXM-specific i-DMRs likely represent a separate class of relevant subpopulation methylation events.

### Given an appropriate gene signature, DXM accurately solves prevalence of underlying cell-types in heterogeneous mixtures

We next sought to determine whether DXM could be used to accurately estimate the prevalence of allelic subpopulations. The analogous problem for calling genetic subclones is based on the general assumption that genomic sequencing analysis of a sample containing a subpopulation with 20% prevalence should yield a peak in the distribution of variant allele frequencies (VAFs) around 20%, which reflects the mutations associated with the subclone. Similarly, if there are CpGs that only are methylated in a certain subclone and the subclone comprises 20% of the sample, the expected fractional methylation would be around 20% methylation. However, in a 20:80 mixture of GCBs and monocytes, we found that the methylation distribution of all CpGs does not have a peak near 20% (Figure 4a), which would make absolute prevalence determination difficult. Considering only CpGs that were found in i-DMRs (Figure 4b) or DMRs detected between GCB cells and monocytes by DSS (Figure 4c) improves the distribution. This suggests that absolute prevalence detection is feasible if an appropriate gene signature is supplied to DXM. We thus defined a gene signature based on DMRs between GCB cells and monocytes from a different set of samples (see methods). Using this signature, DXM predictions have very high agreement with expected prevalence when the expected prevalence of the minor subpopulation is between 15-40% (Figure 4d).

**Figure 4.**
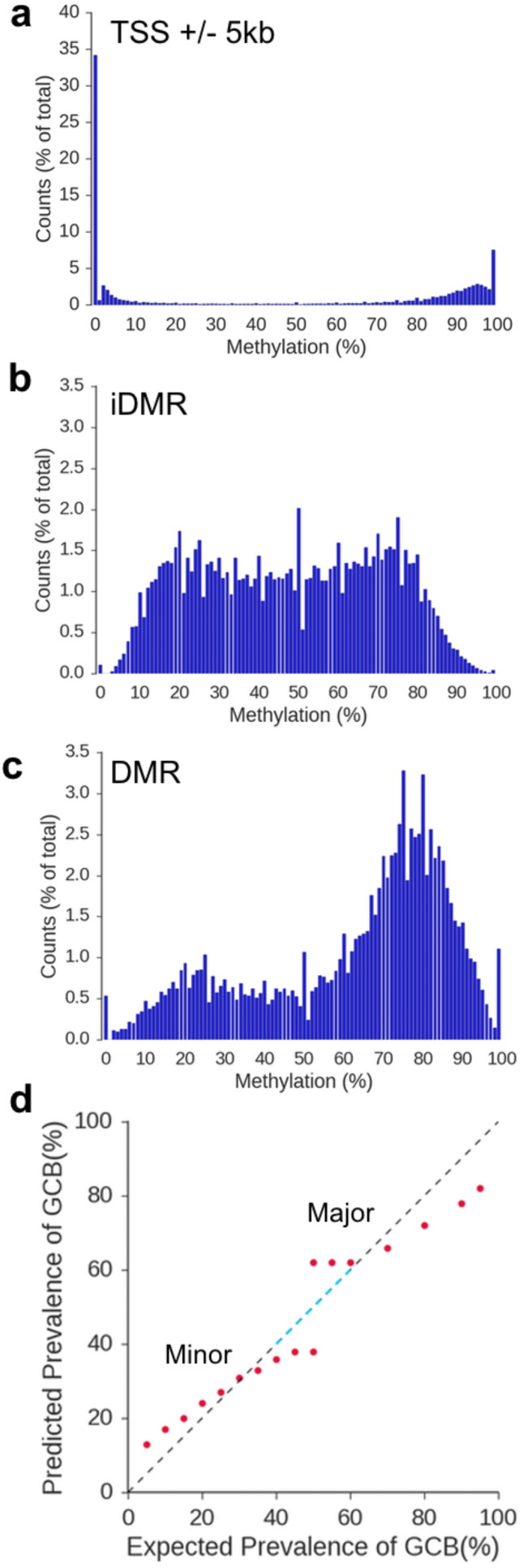
DXM accurately measures the cell prevalence. **a-c)** Methylation distributions of the TSS ± 5kb window (**a**), all iDMRs (**b**), and all DMRs (**c**) from a 55x coverage simulated mixture of monocytes and GCBs (20:80). Expected peaks at ~20% and ~80% percent methylation, which can be used to infer a 20% subpopulation, partially appear using iDMRs and fully appear using DMRs. **d)** Estimated prevalences of GCB cells in GCB:monocyte mixtures (55x coverage) using a GCB-monocyte gene signature. Dashed blue line indicates a perfect estimate.

### DXM predictions for more than two subpopulations

We further sought to evaluate DXM performance for mixtures with more than two subpopulations. Though we found few examples of gene promoters with more than two distinct methylation profiles among cell-types (e.g. a DMR between each pair of cells at that locus, Figure 1h), we applied DXM to a 55x simulated mixture of CD4^+^T:GCB:monocytes (10:25:65) (Supplemental Figure 12). The 496 genes in this mixture showed enrichment in several immune pathways, which is expected from ontology analysis. DXM identified multiple methylation profiles at 434 of the 490 (88.5%) promoters expected to have three methylation profiles, but 342 (79%) of these promoters were solved with only two profiles instead of three (Supplemental Figure 12b). This suggests that in those 342 promoters, DXM identified subpopulation methylation differences but could not resolve them into 3 distinct profiles. For these promoters, DXM had high accuracy for both the major (95.9%) and minor (87.4%) profiles (Supplemental Figure 12c). This accuracy resembles what was seen for k=2 simulations in Figure 2. For the 57 promoters solved with three methylation profiles, DXM had high accuracy for the first (96.0%), second (87.0%), and third (77.6%) profile (Supplemental Figure 12d). Taken together, DXM can solve for more than two subpopulation methylation profiles.

### DXM outperforms existing bisulfite sequencing deconvolution algorithms

We compared DXM with methylPurify(19), an expectation maximization-based approach developed to separate normal and tumor cell profiles from a heterogenous mixture. We generated a 20x coverage simulated mixture of HMEC:HCC1954 cells (35:65) and applied DXM and methylPurify to deconvolve the sample. Both methylPurify and DXM accurately solved methylation profiles of the underlying cell types (methylPurify 98.9%, DXM 97.8%). DXM identified i-DMRs at 58% (15,009/26078) of the most informative bins (mib) predicted by methylPurify, while regions specific to either method tended to cover fewer CpGs (p < 0.001, Cohen’s d>0.8, Supplemental Figure 13). The methylation of partially methylated CpGs in methylPurify mib were also lower than for regions identified by both methods (p < 0.001, Cohen’s d = 4.5). One important difference between the approaches is that since methylPurify uses a binning scheme for methylation changes, it cannot distinguish any methylation changes occurring within a bin (example in Figure 5a).

**Figure 5.**
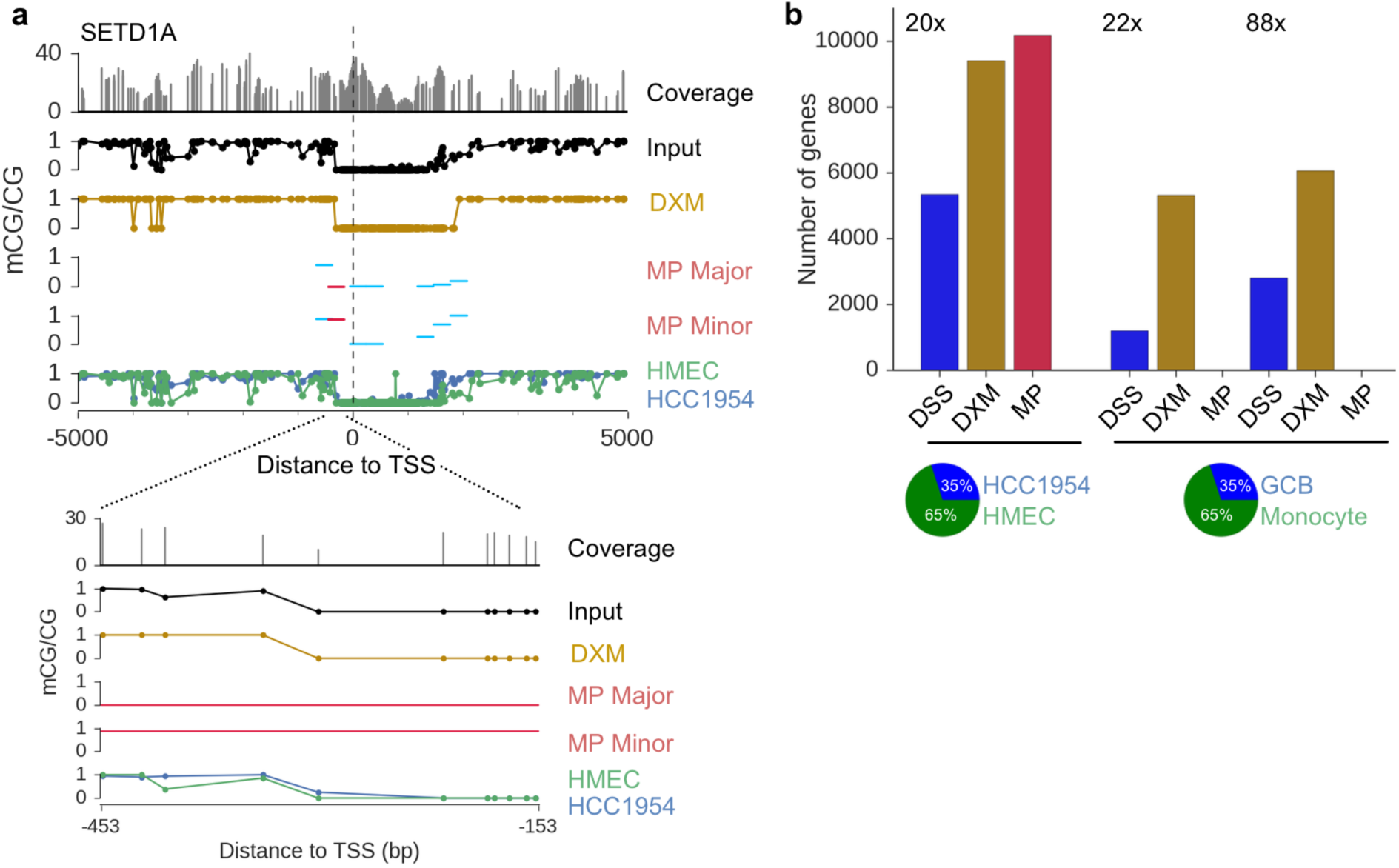
DXM outperforms existing methods. **a**) methylPurify (MP) and DXM outputs for SETD1A in a 20x coverage HMEC-HCC1954 mixture (35:65). Both all (sky blue) and most informative (crimson) bins as determined by MP are depicted. Despite agreement between HMEC and HCC1954 profiles, MP separates this region into two profiles (likely due to the fixed bin size). DXM, however, predicts only one profile. **b**) Number of gene promoters with differential methylation detected by DSS (DMRs), DXM (iDMRs), and MP (two distinct profiles, defined as having at least one bin with >30% methylation difference) across several mixtures. MP fails to identify any promoters with two distinct profiles in GCB-monocyte mixtures at either 22x or 88x coverage.

To our surprise, methylPurify did not perform as well on a 22x coverage simulated mixture of GCBs and monocytes (35:65). It found zero methylation profiles in the mixture for the TSS±5kb window (Figure 5b), in stark contrast with the 1,203 DMRs expected by DSS and 5,319 i-DMRs found by DXM. Additionally, methylPurify estimated the prevalence of the components as 11.5% and 88.5%, while DXM estimated them as 32.5% and 67.5% using a gene signature based on DMRs between GCBs and monocytes as in Figure 4d. Increasing sequencing coverage (88x) did not improve the number of methylation profiles methylPurify found in our window and resulted in a worse prevalence estimate of 5.5% and 94.5%. By comparison, increased coverage leads to DXM prevalence estimates of 33.5% and 66.5% Taken together, we conclude that DXM performs better than methylPurify.

### DXM predicts subpopulation differences in methylation in primary DLBCL samples

We next experimentally validated DXM predictions from Agilent Methyl-Seq analysis of four cryopreserved DLBCLs (sample characteristics are in Supplemental Table 2). Agilent Methyl-Seq is a bisulfite sequencing approach that enriches for gene promoter regions using capture probes similar to exome capture techniques(43). We obtained an average of >54x coverage across ~4.8 million CpGs for all samples, including >58x coverage for >2 million out of ~3.5 million CpGs in the 10kb region around the TSS (Supplemental Table 3). CGIs and the region at the TSS were hypermethylated in DLBCLs relative to normal cell types, while proximal regions were hypomethylated as has been observed in prior DLBCL studies(5) and are typical in most cancers (Figure 6a,b).

**Figure 6.**
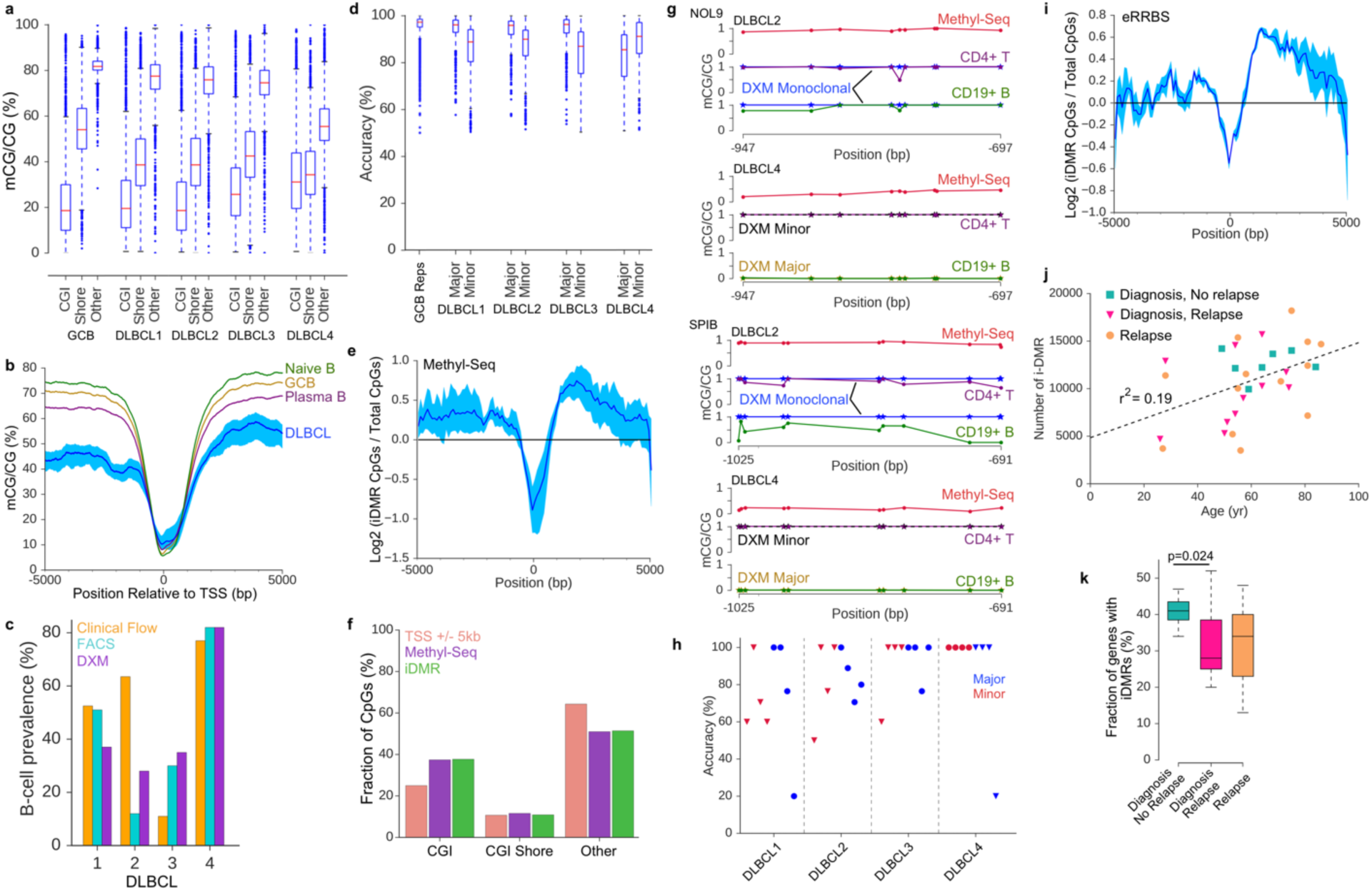
Experimental validation of DXM predictions in DLBCLs. **a)** Distribution of methylation for CpGs in CGIs, CGI shores, or other regions for four DLBCLs profiled by Agilent Methyl-Seq. **b)** Meta-gene analysis (with respect to TSS) of DLBCLs and reference cell types show hypermethylation at and hypomethylation away from the TSS. **c)** Sample prevalence measured by clinical flow (orange), FACS (light blue), and DXM (purple). **d)** Accuracy of DXM predictions for each sample at 2,454 gene promoters with DMRs identified between normal CD4^+^ T-cells and GCB cells. **e)** i-DMRs are enriched 500bp-3kb downstream of the TSS (shading indicates minimum and maximum value across all four samples). **f)** i-DMRs are not enriched in CGIs, CGI shores or other genomic regions relative to the distribution of CpGs covered by Methyl-Seq (p=1.0, Chi-square) **g)** Targeted bisulfite sequencing results for NOL9 and SPIB for two DLBCLs. All results are in Supplemental Figure 13. Methyl-Seq input data (red), DXM predictions (monoclonal blue, major profile gold, minor profile black dash), sorted CD4^+^ T-cells (purple), sorted CD19^+^ B-cells (green). **h)** Accuracy of DXM predicted profiles in a targeted bisulfite sequencing experiment for four samples across four loci (NOL9, SPIB, CD22, BCL2L1) for sorted CD19^+^ B-cells and CD4^+^ T-cells. Blue = major subpopulation, Red= minor subpopulation, triangle = B-cell, circle = T-cell. **i)** i-DMRs are enriched 500bp-3kb downstream of the TSS (shading indicates minimum and maximum value across all 31 samples(5). **j)** Number of i-DMR detected in 31 DLBCLs with respect to patient age (r^2^ = 0.19). **k)** Fraction of gene promoters with an i-DMR for DLBCLs diagnosis (blue-green, n=11), for diagnosis without future relapse (pink, n=7), and at relapse (orange, n=13).

After running DXM on each DLBCL, we first compared DXM-solved estimates (using a gene signature from normal CD4^+^T and GCB cells as above) of the prevalence of T-cell and B-cells, the expected major underlying cell types for DLBCL, to prevalences estimated from clinical flow and FACS. Clinical flow cytometry for CD3+ T-cells and CD19^+^ B-cells was performed at the time of biopsy, while FACS was used to isolate and quantify viable (7-AAD-) CD4^+^ T-cells and CD19^+^ B-cells from cryopreserved samples. We found good agreement between DXM-estimated prevalence and FACS results (on average, fractional prevalence estimates were within 0.0875 of each other, B-cells shown in Figure 6c). Clinical flow data was consistent with FACS and DXM for all but one DLBCL sample (DLBCL 2). DXM results likely are more consistent with FACS sorting because clinical flow could have been obtained from a different region of the biopsy, and DLBCL lymph nodes are known to be heterogeneous.

Next, we estimated the accuracy of DXM subpopulation methylation profiles for the gene signature between normal CD4^+^ T-cells and normal GCB-cells as in Figure 4. For these 2,454 genes, DXM identified differential methylation in 901 - 1,141 promoters with an average accuracy of 91% for major subpopulations and 86% for minor subpopulations, which approaches the levels observed during benchmarking (Figure 6d, Figure 2b). The clear outlier is DLBCL 4, which has predominantly more B-cells than T-cells (Figure 6c) and shows greater CGI hypermethylation. Thus, the major population of DLBCL 4 is likely comprised of malignant B-cells, whose methylation profiles differ substantially from the normal GCBs used as a reference to estimate the accuracy, causing an artificially low estimated accuracy. For each sample, DXM found 3,785 - 6,790 gene promoters with at least one i-DMR (Supplemental Table 3). While i-DMRs were not enriched in CGIs or CGI shores, we did observe an enrichment of i-DMRs in the region located 500-3,000 bp downstream of the TSS (Figure 6e,f), which has been found to strongly associate with expression changes(25).

To experimentally validate DXM profile predictions, we conducted targeted bisulfite sequencing in sorted CD4^+^ T-cells and CD19^+^ B-cells from each sample. We selected four loci that regulate B-cell specific function (SPIB (44), CD22 (45)) or are commonly mutated in DLBCL (BCL2L1 (46), NOL9 (47)) and for which DXM predicted an i-DMR in at least one but not all samples. DXM predicted the correct number of underlying methylation profiles (one or two) across each locus and each patient (example in Figure 6g, data for all 16 cases found in Supplemental Figure 14). Additionally, DXM predictions for subpopulation methylation profiles are recapitulated in sorted cell types with an average accuracy of 84.7% (Figure 6h).

As proof-of-concept, we applied DXM to a publicly available cohort (Cohort 1 in (5)) of 31 DLBCLs (11 paired diagnosis-relapse, 1 of which had 3 relapses, 7 with no relapse) profiled by enhanced reduced representation bisulfite sequencing (eRRBS). These samples exhibit a similar gain of methylation at the TSS and loss of methylation outside the 2 kb surrounding the TSS (Supplemental Figure 15a). DXM identified 3,708 - 18,200 i-DMRs (average of 10,826) for these samples (Supplemental Table 4). i-DMRs showed a similar enrichment 500 - 3,000 bp downstream of the TSS (Figure 6i) and lack of enrichment at CGI and CGI shores (Supplemental Figure 15b), as seen with our DLBCLs (Figure 6). We observed a weak correlation between age and the number of i-DMRs detected for each sample (Figure 6j, r2 = 0.19). This is suggestive that some i-DMRs may be caused by age-associated drift in DNA methylation patterns(39). Additionally, we found that patients presenting with fewer gene promoters with i-DMRs at diagnosis had higher rates of relapse (Figure 6k).

Since an i-DMR represents a gene with multiple methylation profiles, it is possible that the number of genes with i-DMRs in a sample serves as a proxy of the subclonal heterogeneity. This suggests that at diagnosis, samples with fewer i-DMRs, which are less heterogeneous, might represent a more clonal disease, which could be more aggressive and more likely to relapse. Since relapse samples go through a bottleneck where many of the subclones in the diagnosis sample are lost due to treatment, as expected there was no clear relationship between the number of i-DMRs at diagnosis and at relapse. Although these findings, which are based on 18 patients, need to be validated in a larger cohort, they illustrate how DXM could be applied to analyze subpopulation methylation profiles in cancer samples.

## DISCUSSION

We present DXM, a computational method to deconvolve methylation sequencing data from a heterogeneous sample into its major allelic subpopulations, their relative prevalence, and their associated methylation profiles. Importantly, DXM does not require explicit prior knowledge of the expected cell types or number of subpopulations to consider. When using DXM, one consideration is that DXM solves for allelic profiles, which we then interpret as corresponding to unique subpopulations. As such, care must be taken in interpreting DXM results, or those from any deconvolution strategy, in regions with allele-specific methylation or copy number variants. An additional consideration is that DXM does not consider 5-hydroxymethylation (5hmC), which may impact interpreting results in samples with high levels of 5hmC, such as neuronal cell types(48).

If given an appropriate set of regions that are known to show variation in the cell types, DXM can accurately measure the allelic fraction of each cell-type. One potential advantage of using DXM to detect cell prevalences is that DNA methylation is highly stable even in cryopreserved samples, especially relative to RNA and protein. Further, malignant cells are often more susceptible to cell death caused by delays related to specimen transport and cryopreservation (Payton, unpublished observations), which could potentially affect clinical flow and FACS. However, additional testing would be required to demonstrate that DXM could provide a more general way to detect cell prevalences that is beyond the scope of this work.

As a pseudo-reference-free method, DXM does not require reference profiles for each cell-type, but can use them to facilitate interpretation. Instead, DXM uses generalized model parameters to identify subpopulation level methylation profiles. However, if reference profiles are available, they can still be useful to interpret DXM results. This includes the methylation distribution of a typical cell-type as well as the autocorrelations of methylation found in those cell types. To ensure that there was no overfitting for these parameters, we used independent datasets for all training and evaluation steps. We have provided a set of general parameters to use with DXM, but it is possible that certain cases, such as a mutation in a DNA methyltransferase (e.g. DNMT3A in an AML patient(49) or treatment with demethylating agents(50)), may warrant a different set of parameters for optimal performance. In particular one must pay attention to tuning of the beta-binomial error model based on the distribution of methylation across the sample. This models how much noise is expected in individual datasets, and substantially affects the smallest allele fraction that can be detected, as well as the accuracy in determining cell prevalence for low prevalence subpopulations.

A major challenge of genomic methylation data analysis is determining what constitutes a biologically meaningful methylation change, or a distinct methylation profile. One such approach is to consider two profiles distinct if they differ in methylation at any individual CpG, such as in methylFlow(31). However, this can yield a large number of possible subpopulation profiles (e.g. 1.7 billion) that may be intractable for interpretation in many studies. DXM takes an alternative approach and regularizes small differences in methylation between two profiles to report a small number of smoothed profiles. This interpretation is used frequently for imprinting control regions, where partially methylated alleles are considered as biological noise, or drift from a true pattern rather than a functionally different state. A similar assumption underlies DMR-callers. The differences between DSS-called DMRs and i-DMRs in artificial mixtures are similar to those observed between different DMR-callers(51). DMR-callers rely on several important user-defined parameters such as the minimum length, minimum number of CpGs, or minimum difference in methylation between samples, that can severely affect the number and type of DMRs in an analysis(30). Despite this, DMR callers are incredibly useful and are part of every common workflow for DNA methylation analysis.

One surprising result from this analysis is that even sorted cell types have a substantial number of intrinsically intermediate methylated regions, or genomic intervals that are not fully methylated, unmethylated or imprinted. The origin of this observation is not completely understood but likely includes several previously described phenomena including lowly methylated regions (LMRs) and partially methylated domains (PMDs). Intermediate methylation states can be both cell-type specific and conserved (52). Our results show that even in sorted cell types there is extensive intermediate methylation that complicate deconvolution efforts, particularly reference-free cell prevalence determination. This suggests that there are fewer CpGs whose methylation is “cell-type specific” as compared to CpGs with intermediate methylation levels in an individual cell type. Initial reports have demonstrated that LMRs, relatively short non-CpG island DNA segments with low levels of methylation, frequently arise from the binding of transcription factors (53)(54). However, we find near equal levels of highly and lowly methylated DNA methylation in i-DMRs indicating that while LMRs could contribute to the i-DMRS we observe in individual cell types, they cannot fully explain them. PMDs are broad regions of lower methylation, commonly found in cancers and to a limited degree in somatic cell types (55). Cell-type specific PMDs are predicted by CpG density in addition to late replicating/lamina associated domains(56). Given the lack of association of i-DMRs with CpG-density, like LMRs they could contribute to the i-DMRs, but are also unlikely to explain them all.

In summary, here we have generated a side-by-side comparison of DXM to existing allelic subpopulation methylation analysis techniques and demonstrated superior performance in methylation profiles detected. Further, we have validated that in DLBCL samples, subpopulation methylation profiles predicted by DXM are recapitulated in relevant sorted cell subpopulations. As proof-of-concept, we have used DXM to identify subpopulation methylation profiles in DLBCL that may correlate with patient prognosis. We expect that DXM will have high applicability and utility for the analysis of subpopulation methylation in any heterogeneous sample.

## AVAILABILITY

DXM is available on github at https://github.com/edwardslab-wustl/dxm under the GNU GPL License. DXM was written in Python3 and was tested on CentOS7, though it should work on any *nix system. For DLBCL samples, DXM on average used 300Mb memory and processed a sample in less than two hours, using one CPU. A docker container for dxm can be found at https://hub.docker.com/repository/docker/jedwardswustl/dxm.

## SUPPLEMENTARY DATA

Supplementary Data are available online.

## ACKNOWLEDGEMENT

We thank Yu Sun for his input into early versions of the deconvolution algorithm. We thank the Genome Technology Access Center for help with genomic sequencing and the Flow Cytometry Core in the Department of Pathology and Immunology for help with cell sorting experiments. We thank the Washington University School of Medicine (WUSM) Lymphoma Banking Program, Dept. of Medicine, Division of Oncology and N. Bartlett for lymphoma biopsies.

