## Supplementary Figures and Tables for "Determining subpopulation methylation profiles from bisulfite sequencing data of heterogeneous samples using DXM"

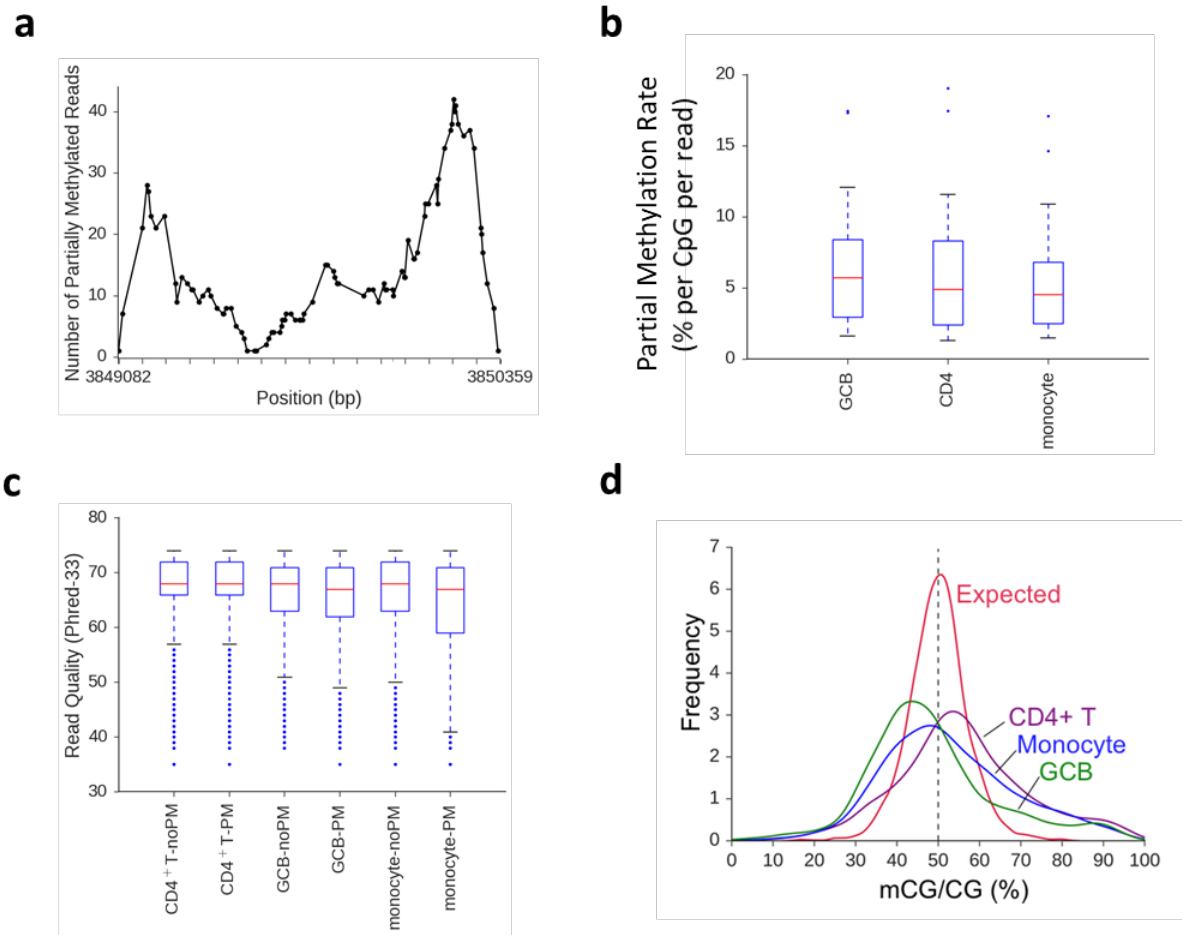


**Supplemental Figure 1. Partially methylated reads span imprinted DMRs, cannot be filtered by read quality, and result in wider distributions of detected methylation. a)** Distribution of partially methylated reads across the FAM50B locus in GCB cells. Reads occur throughout the entire locus (ticks every 80bp). **b)** Average rate (per CpG, per read) of detecting a CpG with partial methylation in imprinted loci (GCB-8.3%, monocyte-7.7%, $CD4^{+}$T-6.9%). **c)** Distribution of read quality for partially methylated reads (PM) or those that are fully methylated or unmethylated (e.g. not partially methylated noPM). The read quality is not substantially different between PM and noPM reads for $CD4^{+}$T cells (d=0.11), GCB cells (d=0.187), and monocytes (d=0.22). d = Cohen's effect size. **d)** Fractional methylation of CpGs in imprinted DMRs for three sorted cell types. Distribution is wider in sorted cell types than expected for 50% methylation (red dotted line). Note that data are for single end sequencing so the number of reads and fragments are equivalent.

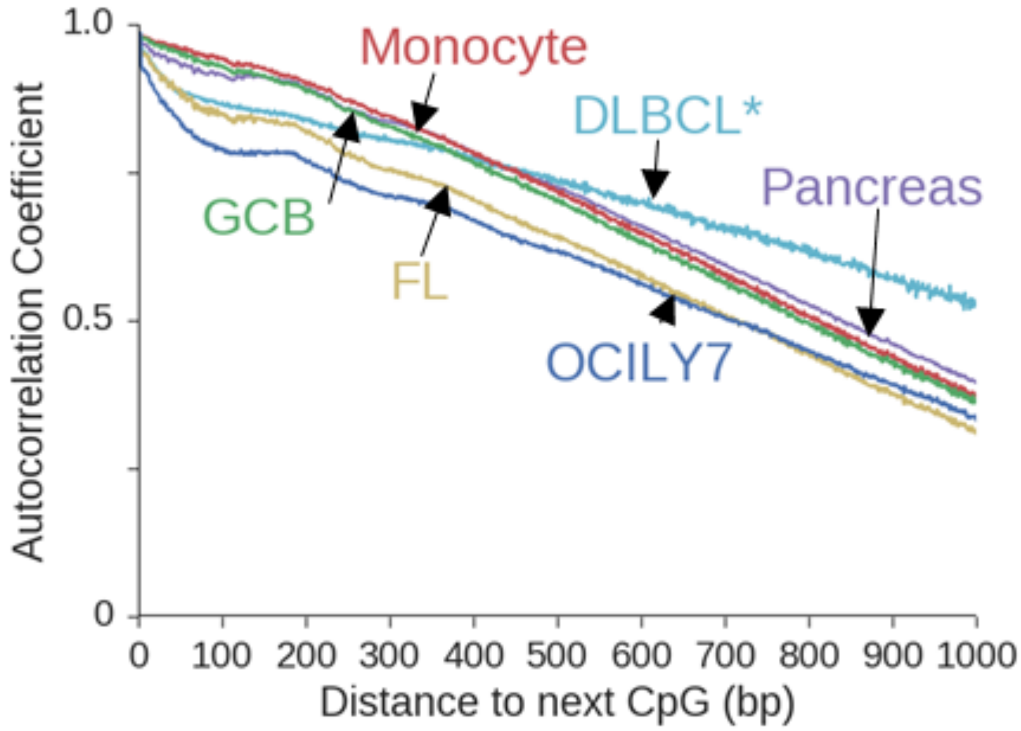


**Supplemental Figure 2. Methylation exhibits high degrees of correlation between CpGs.** Representative samples were selected for Follicular Lymphoma (FL) and DLBCL. *denotes ERRBS, all other samples are WGBS.

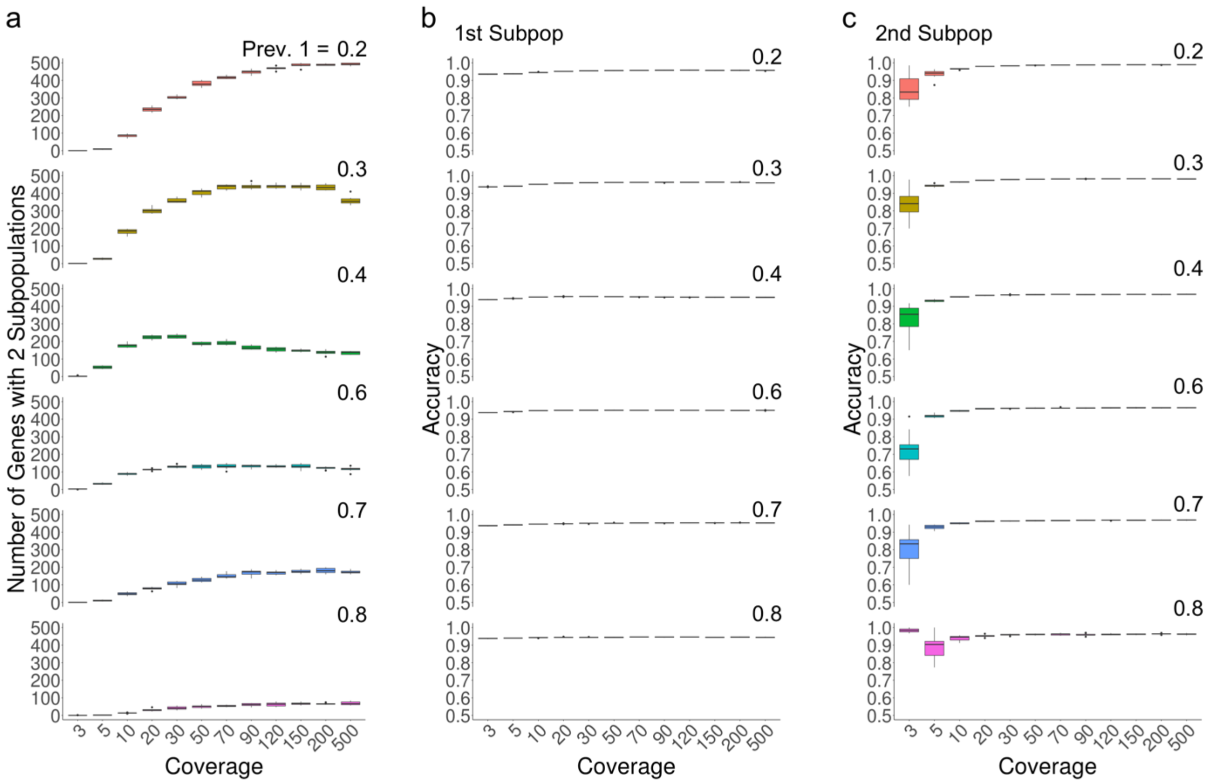


**Supplemental Figure 3. DXM performs well on simulated data.** DXM was used to deconvolve 720 simulated mixtures of HMEC and CD4T cells. Methylation values were binarized prior to simulation. The prevalence of HMEC cells in the mixture (Prev. 1) was varied from 0.2 to 0.8. The coverage was fixed for each simulation, and mixed methylation values were modelled based on binomial sampling. For each coverage and prevalence, 10 simulations of 1000 genes were performed. The total number of genes with two subpopulations is shown (**a**) along with the accuracy for the 1$^{st}$ and 2$^{nd}$ subpopulations (**b** and **c**). DXM only separates out a second subpopulation when the accuracy is high.

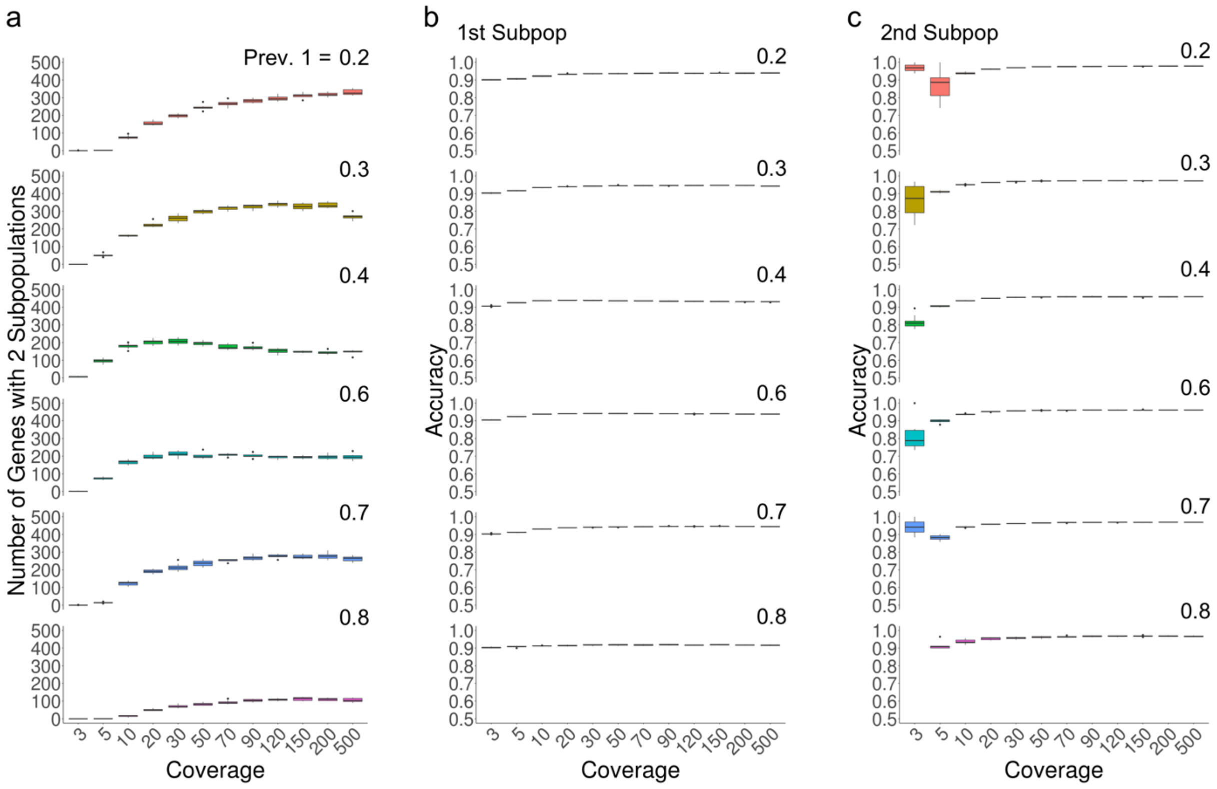


**Supplemental Figure 4. DXM performs well on simulated data.** DXM was used to deconvolve 720 simulated mixtures of HMEC and HCC1954 cells. Methylation values were binarized prior to simulation. The prevalence of HMEC cells in the mixture (Prev. 1) was varied from 0.2 to 0.8. The coverage was fixed for each simulation, and mixed methylation values were modelled based on binomial sampling. For each coverage and prevalence, 10 simulations of 1000 genes were performed. The total number of genes with two subpopulations is shown (**a**) along with the accuracy for the 1$^{st}$ and 2$^{nd}$ subpopulations (**b** and **c**). DXM only separates out a second subpopulation when the accuracy is high.

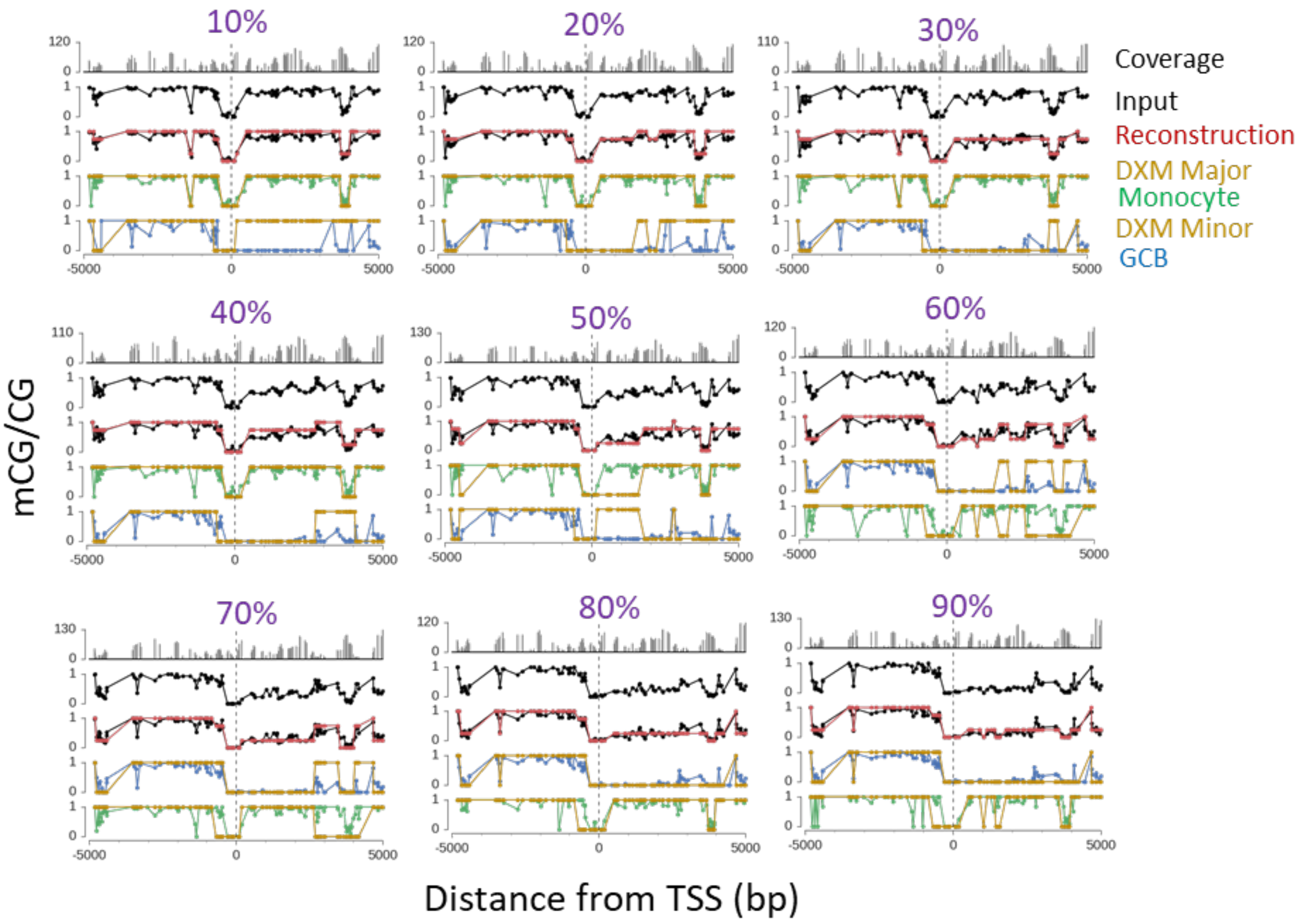


**Supplemental Figure 5. DXM performance is high when one subpopulation is greater than 10% prevalence and when both subpopulations are not at the same prevalence (50%).** DXM solutions for the CD22 gene across GCB:monocyte mixtures with 55x average coverage and varying prevalence. Percentage denotes GCB prevalence.

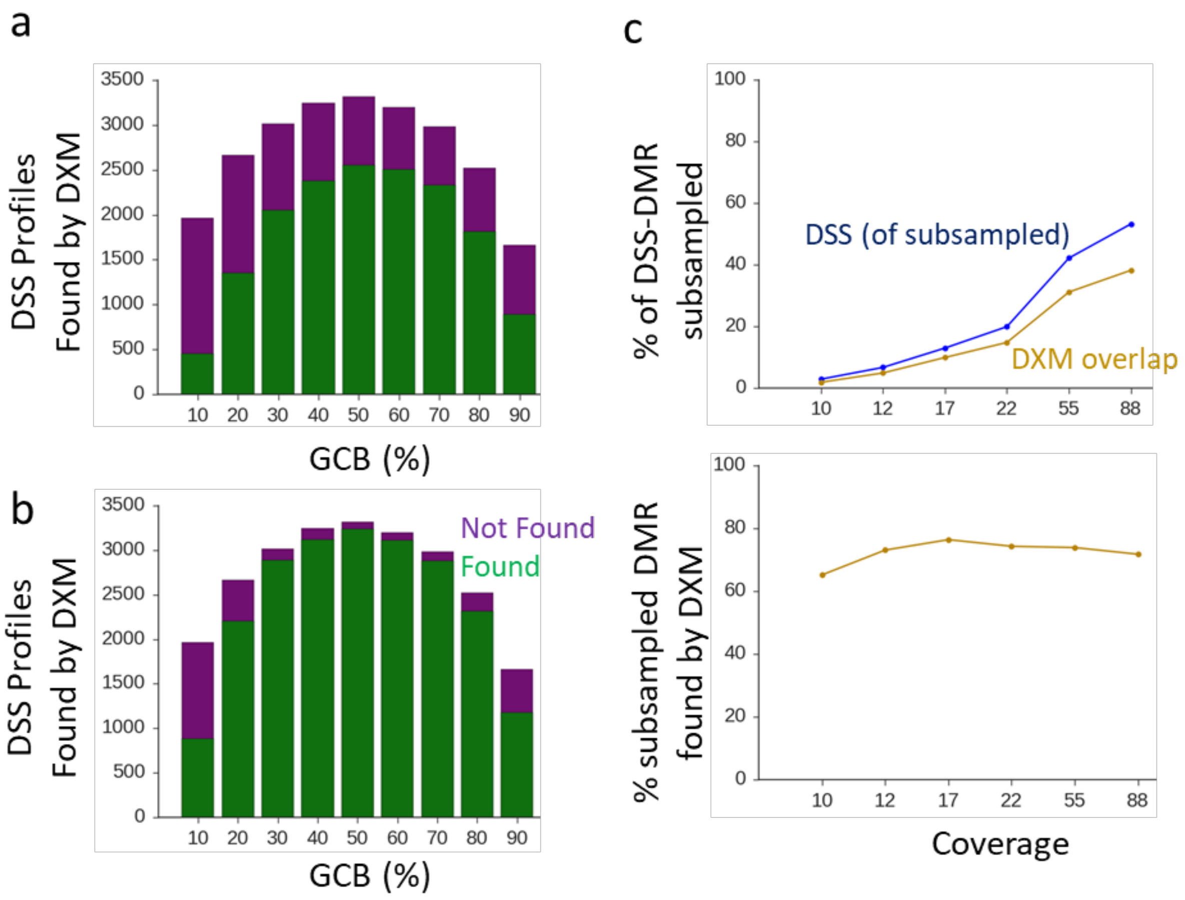


**Supplemental Figure 6. DXM i-DMRs recapitulate DMRs identified by DSS. a)** Number of DSS-identified DMRs that are not found (purple) or found (green) by DXM as i-DMRs. DXM identifies an average of 63.8% of DMRs. **b)** Number of DSS-identified DMRs that are not found (purple) or found (green) by DXM with a more specific domain (target region). Given specific DMR-locations to consider, DXM identifies an average of 85.9% of DMRs. **c)** Number of total DMRs and DMRs that intersect i-DMR identified in subsampled mixtures (35:65 GCB:monocyte) relative to the number found between reference cell types. As coverage increases, the numbers of both DMRs and i-DMRs increase, but the relative number of DMRs recapitulated by DXM does not.

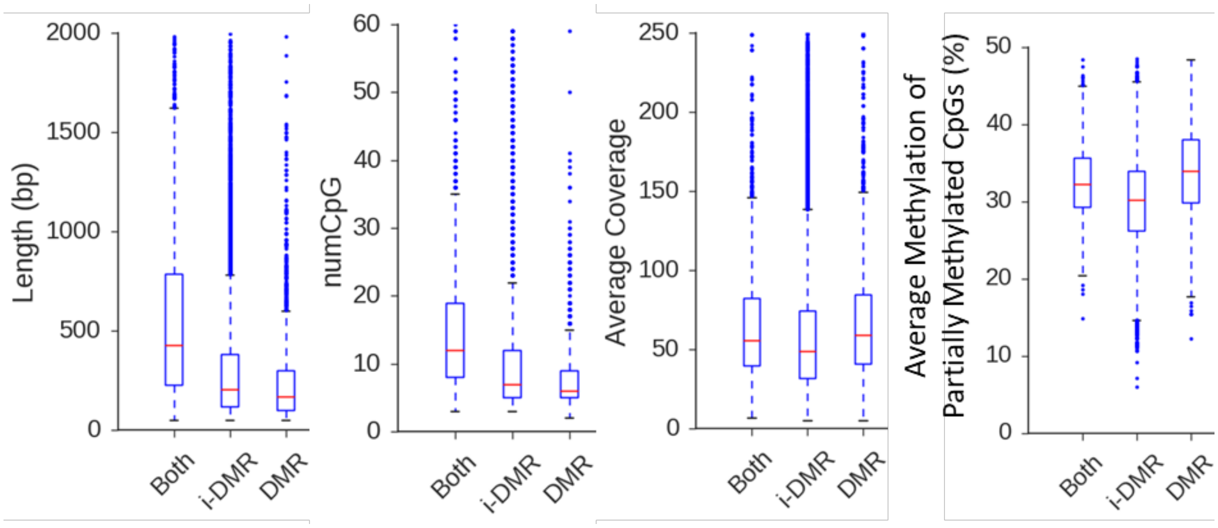


**Supplemental Figure 7. DXM i-DMRs and DSS DMRs that overlap are longer and tend to have more CpGs**. In a 30:70 simulated mixture of GCB:monocytes, regions called by both DXM and DSS (both) tend to be shorter and involve fewer CpG than those called only by DXM (i-DMR) or DSS (DMR). For length, both vs i-DMR ($p<0.001$ ANOVA with Tukey's posthoc, Cohen's $d=0.81$) and both vs DMR ($p<0.001$, $d=0.79$). For number of CpG, both vs i-DMR ($p<0.001$, $d=0.39$), both vs DMR ($p<0.001$, $d=0.79$). There is no difference in average coverage between Both and DMR, and there is a small difference between Both and i-DMR ($p<0.001$, $d=0.16$) and between Both and DMR ($p<0.001,d=0.21$) for average coverage. For average methylation at partially methylated reads, there is a larger difference between Both and i-DMR ($p<0.001$, $d=0.44$) and between Both and DMR ($p<0.001$, $d=0.66$). There is a small difference between i-DMR and DMR for average methylation ($p<0.001$, $d=0.25$).

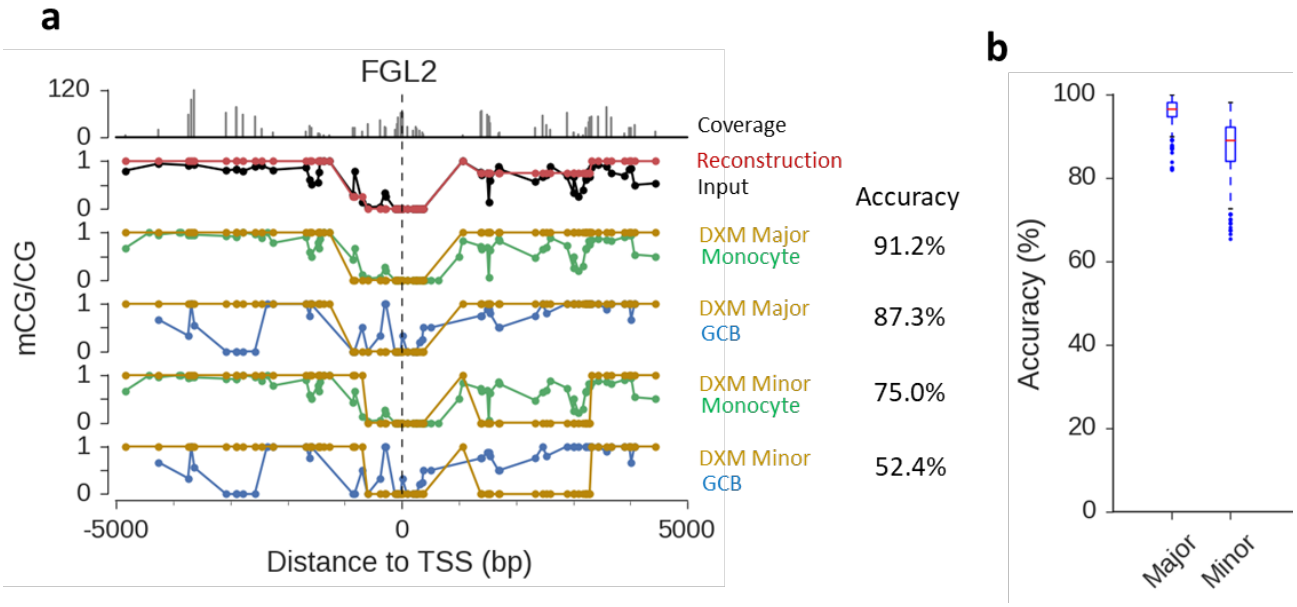


**Supplemental Figure 8. DXM solutions for genes with difficult cell-type assignment. a)** In a 10:90 simulated mixture of GCB:monocytes, DXM identifies two profiles at FGL2, both of which are closest to reference monocytes. The accuracy of each profile to each potential reference is shown on the right. **b)** Accuracy of DXM major and minor profiles for genes where both profiles share the same closest reference in a 10:90 mixture of GCB:monocytes.

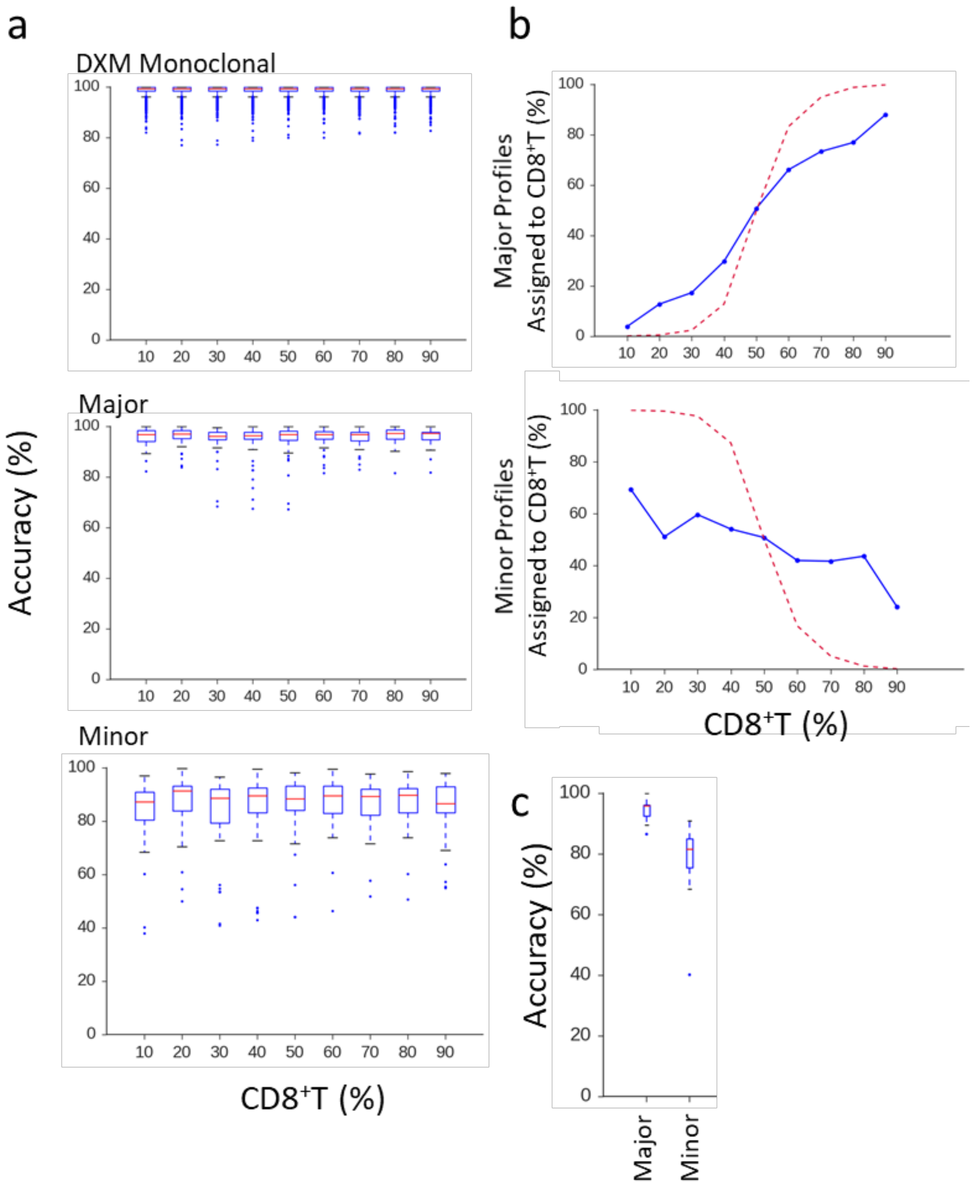


**Supplemental Figure 9. DXM accurately deconvolves major and minor subpopulation profiles from 30x mixtures of CD4$^+$T:CD8$^+$T. a)** Accuracy of methylation profiles to reference cell types for DXM monoclonal genes, or for the major and minor profiles. **b)** Assignment of major (top) and minor (bottom) profiles to cell-types. CD4$^+$T and CD8$^+$T cells are closer in lineage (435 promoters with DMR) than GCB:monocytes (2,988 promoters with DMR), leading to more difficult assignment. **c)** Accuracy for cases where there both major and minor profiles solved by DXM are assigned to the same cell type (see example in Supplemental Figure 8a)

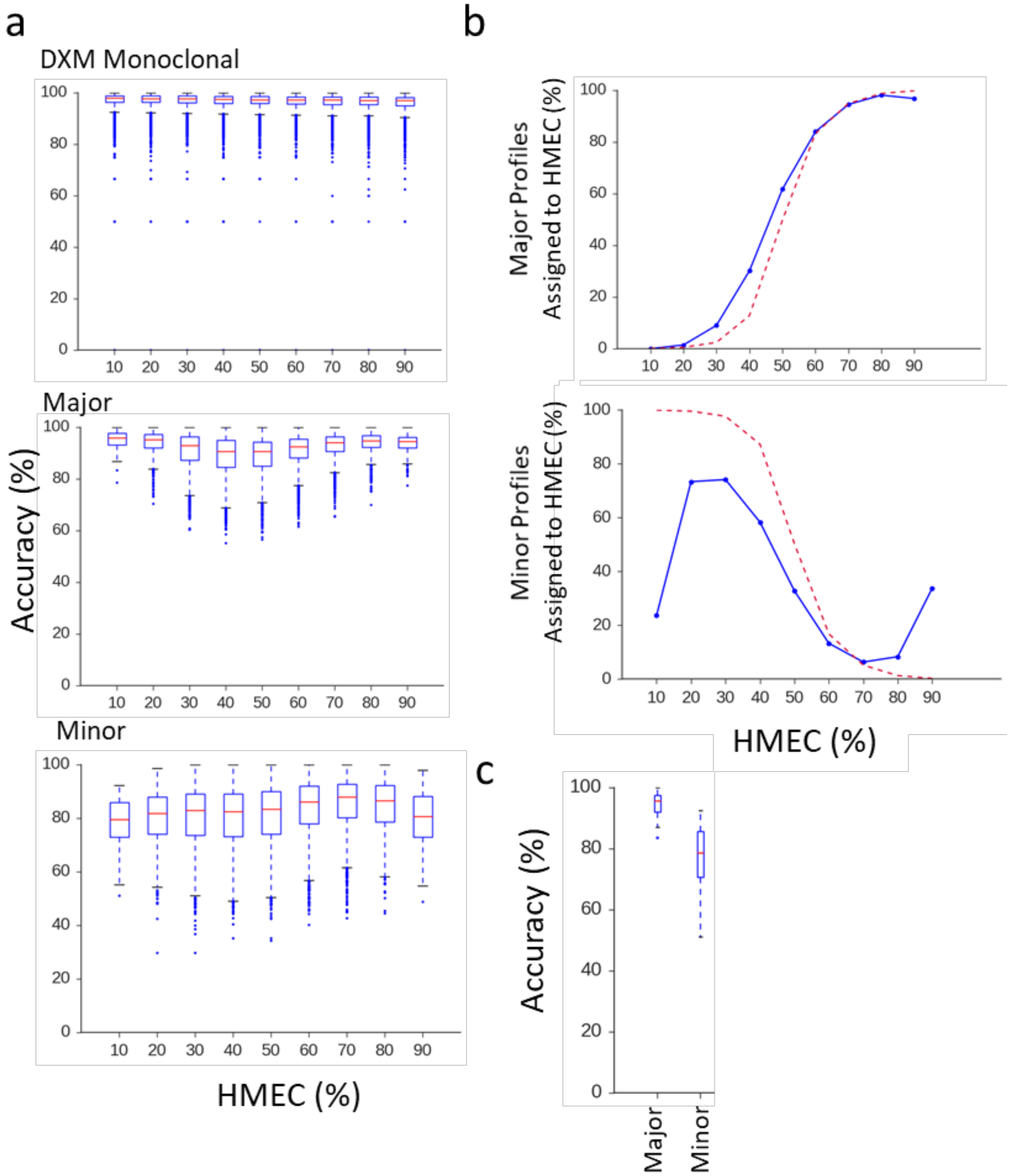


**Supplemental Figure 10. DXM accurately deconvolves major and minor subpopulation profiles from 20x mixtures of HMEC:HCC1954. a)** Accuracy of methylation profiles to reference cell types for genes with a single profile, or the major and minor methylation profile. **b)** Assignment of major (top) and minor (bottom) profiles to cell-types. **c)** Accuracy for cases where there both major and minor profiles solved by DXM are assigned to the same cell type (see example in Supplemental Figure 8a*).*

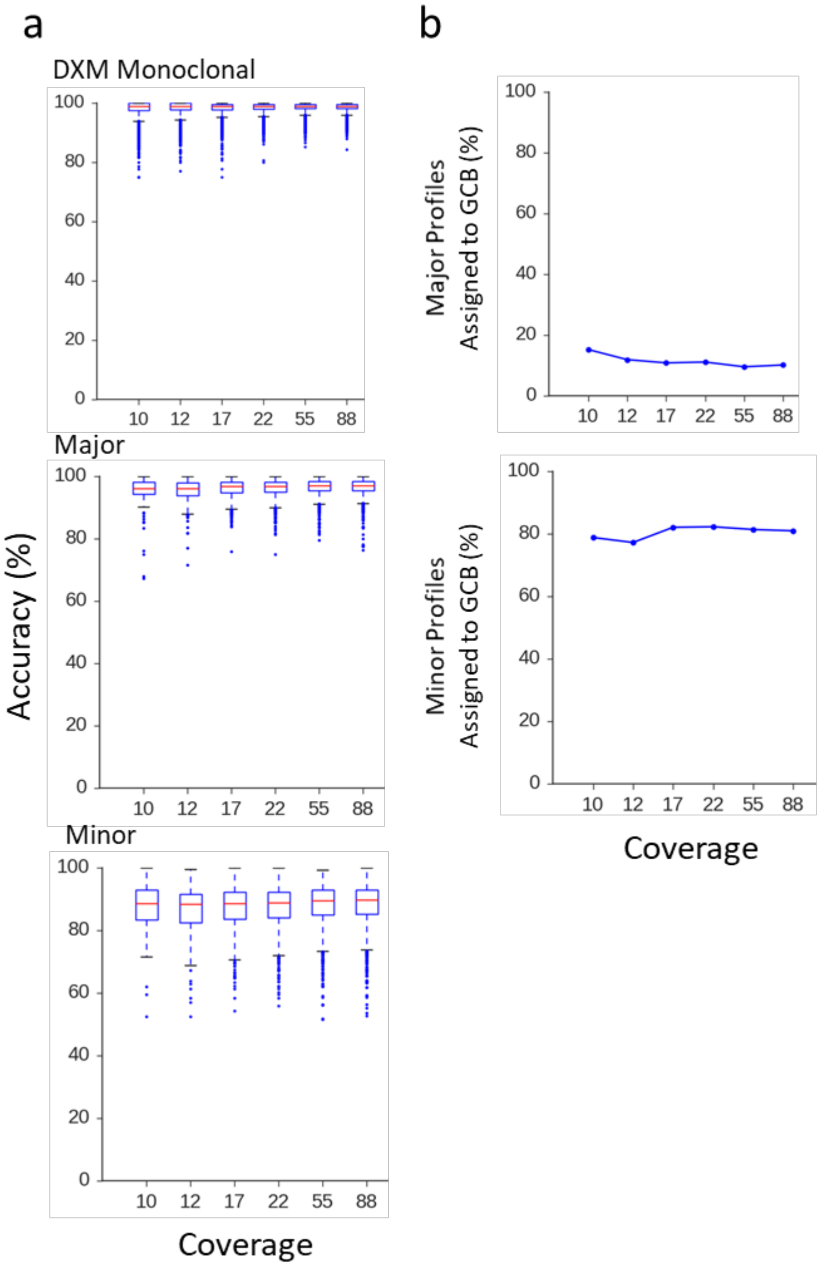


**Supplemental Figure 11. Sequencing coverage from 10x-80x does not affect accuracy of reconstructed profiles or assignment of profiles to cell-types**. DXM i-DMRs were defined from a GCB-monocyte mixture with fixed prevalence (35:65) and varying average global coverage. a**)** DXM accuracy for genes with one profile (single), or the major and minor profiles. **b)** Percent of all major (top) or minor (bottom) profiles assigned to GCB for simulated mixtures.

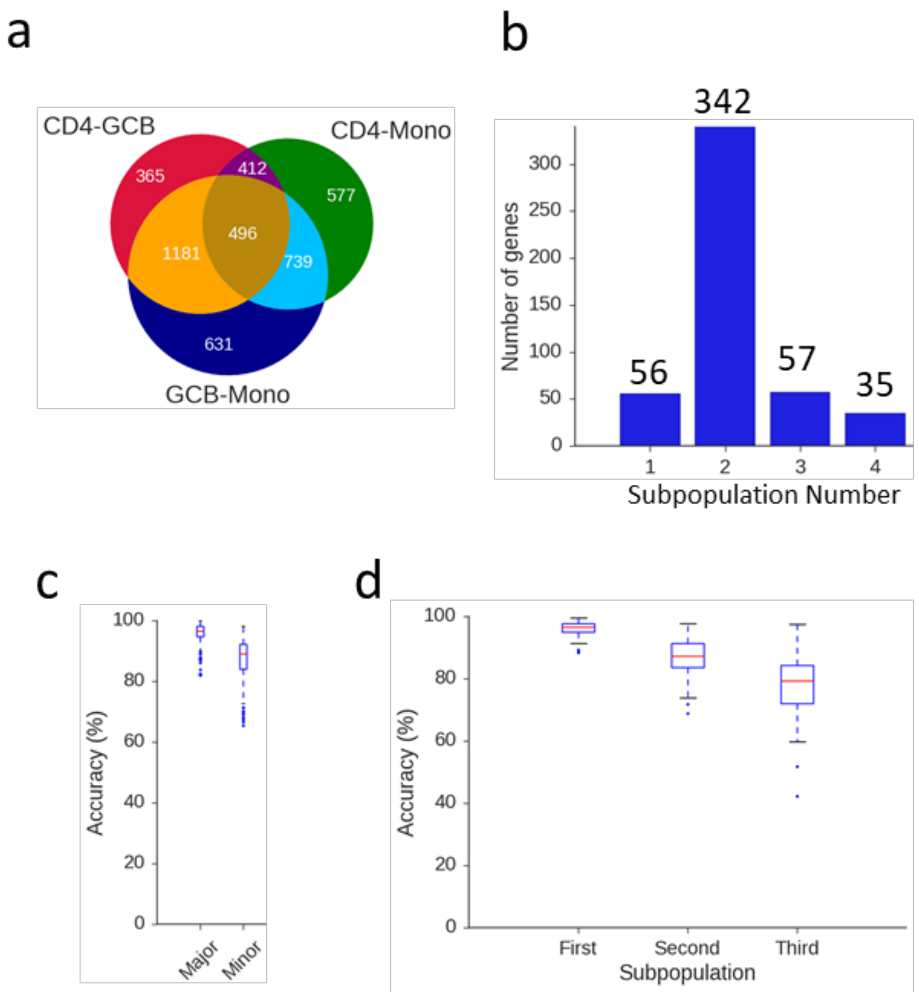


**Supplemental Figure 12. Characterization of DXM performance for more than 2 subpopulations. a)** Venn Diagram for genes with a DMR between at least two cell-types (GCB, CD4T, monocytes). Only 496 genes have a DMR between each pair of cell-types (three expected methylation profiles). **b)** Number of methylation profiles identified by DXM for 490 genes with three expected methylation profiles in a 55x 10:25:65 mixture of CD4T:GCB:monocytes. 6 were not present following subsampling used to generate the mixture. **c)** DXM has high accuracy for major (95.9%) and minor (87.4%) profiles at 342 genes where only two methylation profiles were detected. Performance remains high **d)** DXM has high accuracy of first (96.0%), second (87.0%), and third (77.6%) methylation profiles, ordered by descending prevalence, for the 57 genes where DXM found three distinct profiles.

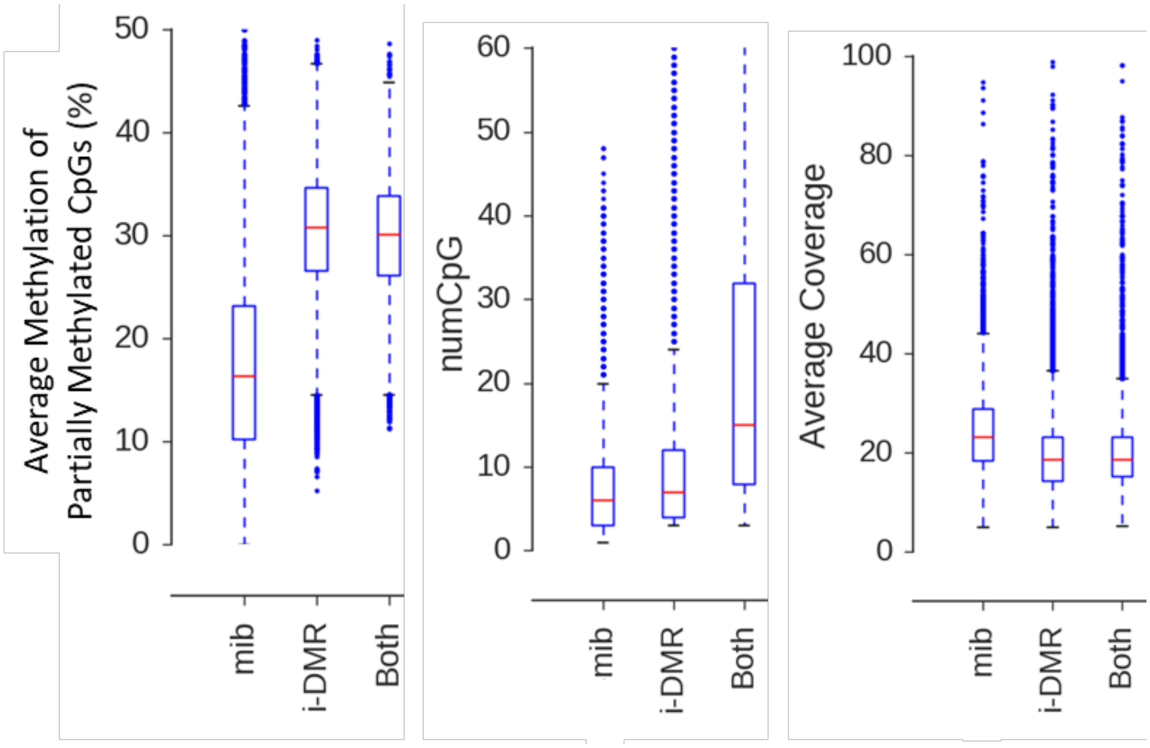


**Supplemental Figure 13. The most informative bins (mib) identified only by MethylPurify tend to have fewer CpG and less pronounced methylation at partially methylated regions than those identified by DXM (i-DMR) or by both methods**. By definition, mibs must be 300bp, whereas DXM i-DMRs can vary in size. For all characteristics, all groups differ from each other ($p < 0.001$, ANOVA with Tukey's posthoc test). There is a substantial effect size (Cohen's d >0.8) for all comparisons except between "Both" and "i-DMR" for partially methylated CpGs (d=0.245) and average coverage (d=0.269).

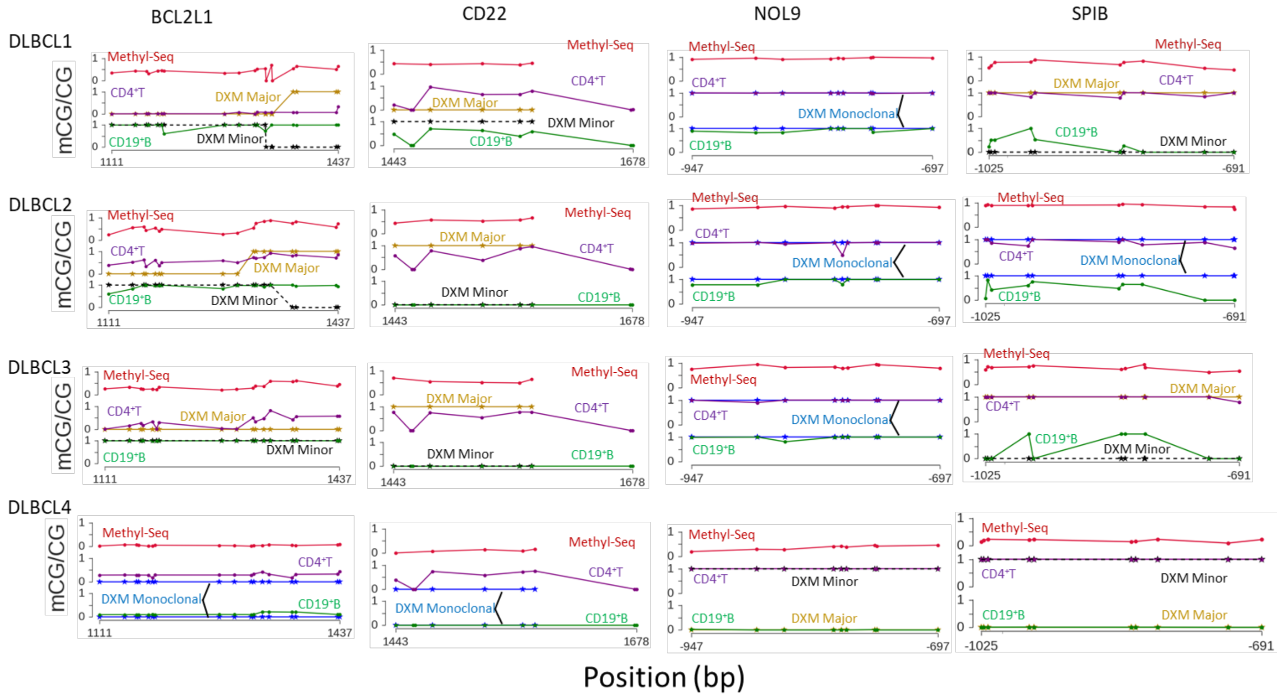


**Supplemental Figure 14. Targeted bisulfite sequencing for four genes in sorted CD4$^{+}$T and CD19$^{+}$B cells from four DLBCL samples.** Each gene had at least one sample predicted to have an i-DMR and at least one without. Legend and colors are the same as Figure 6g.

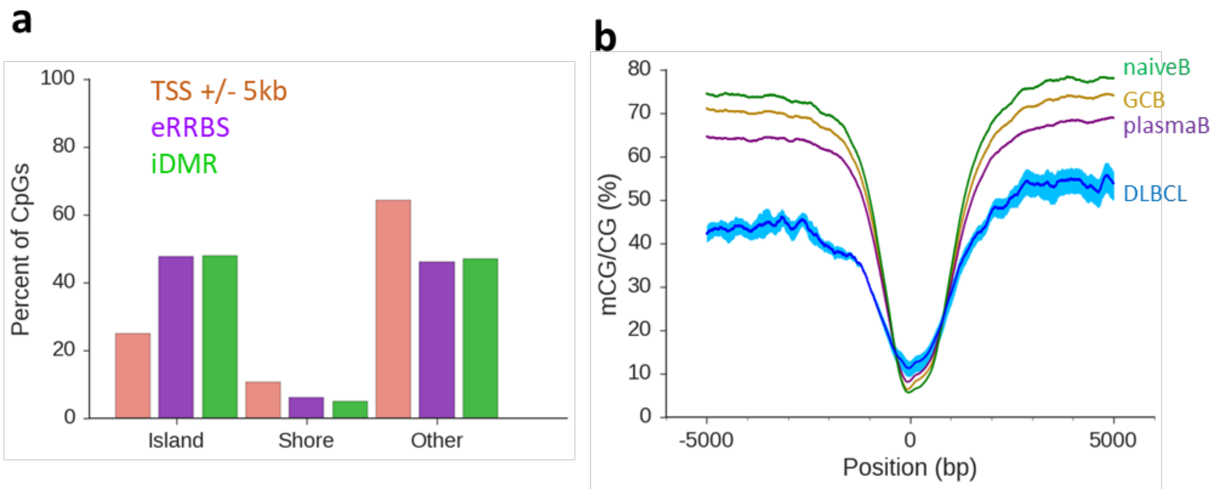


**Supplemental Figure 15. DXM i-DMRs in DLBCLs profiled by ERRBS from Pan et al. 2015, Nature Communications, have similar characteristics as i-DMRs identified in our cohort**. **a)** Location of i-DMRs with respect to CGI, shores, or other regions. **b)** Fractional methylation of DLBCL samples across TSS window. Dark blue denotes the average sample profile, with light blue indicating the maximum and minimum range.

**Supplementary Table 1.** Ontology analysis with DAVID functional annotation tool for genes with an i-DMR found within 9 sorted cell types (all clusters with at least one element with FDR < 0.05).

Annotation Cluster 1: Enrichment Score 42.57

| Category | Term | Count | List Total | Fold Enrichment | FDR |
|---|---|---|---|---|---|
| UP_SEQ_FEATURE | domain:Cadherin 6 | 54 | 796 | 16.39829 | 1.72E-50 |
| UP_SEQ_FEATURE | domain:Cadherin 5 | 54 | 796 | 12.96246 | 6.74E-43 |
| UP_SEQ_FEATURE | domain:Cadherin 3 | 54 | 796 | 12.1523 | 5.49E-41 |
| UP_SEQ_FEATURE | domain:Cadherin 4 | 54 | 796 | 12.1523 | 5.49E-41 |
| UP_SEQ_FEATURE | domain:Cadherin 1 | 54 | 796 | 11.83529 | 3.20E-40 |
| UP_SEQ_FEATURE | domain:Cadherin 2 | 54 | 796 | 11.83529 | 3.20E-40 |
| INTERPRO | IPR020894:Cadherin conserved site | 54 | 762 | 11.63898 | 4.91E-40 |
| INTERPRO | IPR002126:Cadherin | 54 | 762 | 11.1458 | 8.55E-39 |
| INTERPRO | IPR015919:Cadherin-like | 54 | 762 | 10.96004 | 2.55E-38 |
| SMART | SM00112:CA | 54 | 458 | 10.22207 | 1.31E-37 |
| GOTERM_BP_DIRECT | GO:0007156~homophilic cell adhesion via plasma membrane adhesion molecules | 54 | 711 | 8.071783 | 2.35E-30 |

Annotation Cluster 2: Enrichment Score 18.52

| Category | Term | Count | List Total | Fold Enrichment | FDR |
|---|---|---|---|---|---|
| UP_SEQ_FEATURE | repeat:PXXP 5 | 15 | 796 | 25.20477 | 5.20E-16 |
| UP_SEQ_FEATURE | repeat:PXXP 3 | 15 | 796 | 25.20477 | 5.20E-16 |
| UP_SEQ_FEATURE | repeat:PXXP 4 | 15 | 796 | 25.20477 | 5.20E-16 |
| UP_SEQ_FEATURE | repeat:PXXP 1 | 15 | 796 | 25.20477 | 5.20E-16 |
| UP_SEQ_FEATURE | repeat:PXXP 2 | 15 | 796 | 25.20477 | 5.20E-16 |

Annotation Cluster 3: Enrichment Score 5.91

| Category | Term | Count | List Total | Fold Enrichment | FDR |
|---|---|---|---|---|---|
| UP_SEQ_FEATURE | domain:Peptidase S1 | 19 | 796 | 4.393493 | 4.62E-04 |
| INTERPRO | IPR001314:Peptidase S1A, chymotrypsin-type | 19 | 762 | 4.095197 | 0.001224 |
| INTERPRO | IPR001254:Peptidase S1 | 19 | 762 | 3.85631 | 0.003014 |
| SMART | SM00020:Tryp_SPc | 19 | 458 | 3.535693 | 0.007241 |

**Supplementary Table 2.** DLBCL sample characteristics.

| DLBCL | Treatment | Diagnosis | Clinical Flow | IHC | FISH |
|---|---|---|---|---|---|
| 1 | Diagnosis | DLBCL-recurrent, nonGCB | REGION A small cells: CD3: 63%; CD19: 37%; Region B large cells: CD3: 37%, CD19: 68% | CD20+, PAX5+, MUM-1+, BCL6 weak in a subset; CD10- | BCL2, MYC, BCL6- |
| 2 | Relapse | DLBCL-GCB | LN: CD3 24%; CD19 76%; Tonsil: CD19: 49%; CD3: 51% | CD10+, BCL6+, CD5+, BCL2+, CD20+ | None |
| 3 | Diagnosis | DLBCL-GCB | Region A low SS: CD3: 82%, CD19: 15%; Region B high SS:CD19 7% | CD20+, CD10+, PAX5+, BCL2+, BCL6 WEAK | None |
| 4 | Diagnosis | DLBCL-GCB | CD3: 13%, CD19: 77% | CD10+, CD20+ | Igh-BCL2 +, loss of one copy of 8q24 |

**Supplementary Table 3.** Sequencing results and relevant DXM results.

| DLBCL | numCpG | Average Coverage | Non-CpG Conversion (%) | Average Coverage in TSS±5kb window | i-DMR | Genes with i-DMR |
|---|---|---|---|---|---|---|
| 1 | 4.8e6 | 53.2 | 98.9 | 58.7 | 4,746 | 6,758 |
| 2 | 4.8e6 | 53.0 | 98.9 | 59 | 3,785 | 6,087 |
| 3 | 4.9e6 | 54.4 | 98.8 | 60.5 | 6,790 | 9,167 |
| 4 | 4.8e6 | 56.8 | 99.0 | 62.6 | 4,808 | 6,896 |

**Supplementary Table 4**. Number of i-DMR identified in analysis of 31 DLBCL samples (Pan et al., 2015, Nature Communications)**.**

| Sample | i-DMR | Genes with i-DMR |
|---|---|---|
| 1D | 10,113 | 2,961 |
| 2D | 9,002 | 2,885 |
| 3D | 7,341 | 2,044 |
| 4D | 12,907 | 3,827 |
| 5D | 14,564 | 5,182 |
| 6D | 4,700 | 1,603 |
| 7D | 5,300 | 1,664 |
| 8D | 15,703 | 3,899 |
| 9D | 11,738 | 3,309 |
| 10D | 10,276 | 2,912 |
| 11D | 6,477 | 2,269 |
| 1R1 | 12,427 | 2,654 |
| 1R2 | 7,167 | 2,952 |
| 1R3 | 14,929 | 4,041 |
| 2R | 11,543 | 3,448 |
| 3R | 3,506 | 948 |
| 4R | 11,387 | 3,160 |
| 5R | 15,380 | 4,744 |
| 6R | 3,708 | 1,306 |
| 7R | 5,218 | 1,685 |
| 8R | 18,200 | 4,005 |
| 9R | 14,685 | 3,502 |
| 10R | 10,771 | 2,791 |
| 11R | 10,034 | 3,386 |
| NR1 | 13,658 | 4,182 |
| NR2 | 12,236 | 3,850 |
| NR3 | 14,215 | 4,368 |
| NR4 | 12,151 | 3,768 |
| NR5 | 9,952 | 3,111 |
| NR6 | 14,015 | 4,186 |
| NR7 | 12,289 | 3,493 |

**Supplementary Table 5.** Primer information for targeted bisulfite sequencing. All coordinates are hg19.

| Gene | Chr | Strand | Start | End | Primer-1 | Primer-2 |
|---|---|---|---|---|---|---|
| CD22 | 19 | + | 35821466 | 35821822 | AAAAATGTATAGAGTTGGTTAAATAAAA | CATAAACAAATACCCAACAACCTTTA |
| SPIB | 19 | + | 50921140 | 50921537 | GGATTTGGAAGATTAGGAGTAGTT | AATCCCCAAAATCATCACCA |
| BCL2L1 | 20 | - | 30309417 | 30309813 | AGGTTAAAGAAAAGGGATATATAAGGG | ACCACAACAACAATTTAAATACC |
| NOL9 | 1 | - | 6615330 | 6615720 | GGGGTTTGAAGTTAGGAAGTATT | AAAAACAAAAACAAAATCCTATTTA |
